# Carbenoxolone disrupts cell migration by inhibiting the SERCA pump

**DOI:** 10.64898/2026.08.11.743254

**Authors:** Cinthia Sanchez-Rabadan, Belen Calvo, Svitlana Palii, Marie R. Adler, Juan L. Cortes-Muñoz, Christian Conze, Arturo Jiménez-Sánchez, Martin L. Gallegos-Gomez, Ulrike Uhrig, Thomas Schimmang, Jonathan Rojo-Ruiz, Pablo J. Sáez, Maria Teresa Alonso

**Affiliations:** Unidad de Excelencia, Instituto de Biomedicina y Genética Molecular de Valladolid (IBGM), Universidad de Valladolid y Consejo Superior de Investigaciones Científicas (CSIC), Valladolid, Spain; Cell Communication and Migration Laboratory, Institute of Biochemistry and Molecular Cell Biology, Center for Experimental Medicine, University Medical Center Hamburg-Eppendorf, Hamburg, Germany; Bioorganic Chemistry Laboratory (BioChela). Instituto de Química. Universidad Nacional Autónoma de México (UNAM), Mexico City, Mexico; Technology Platform Microscopy and Image Analysis, Leibniz Institute of Virology, Hamburg, Germany; Chemical Biology Core Facility, European Molecular Biology Laboratory (EMBL); Heidelberg, Germany

**Keywords:** Carbenoxolone, gap junctions, calcium, calcium signalling, ER, SERCA, calcium release

## Abstract

Collective cell migration is a fundamental process driving tissue repair, angiogenesis, and vascular homeostasis. This coordinated movement requires both intercellular communication via gap junctions and precise intracellular Ca²⁺ signaling, largely regulated by the sarco(endo)plasmic reticulum Ca^2+^ ATPase (SERCA) pump within the endoplasmic reticulum (ER). Historically, carbenoxolone (CBX)—a synthetic derivative of glycyrrhetinic acid—has been widely utilized as a pharmacological tool to inhibit gap junctions and dissect their role in collective cell motility. However, its molecular specificity remains highly controversial. In the present study, using different cellular models, we found that CBX drastically reduces collective cell migration by a previously undescribed function for CBX: a fast, potent, and reversible inhibition of the SERCA pump, which provokes a passive leak of the luminal ER Ca²⁺ store. Our findings suggest that the effect of CBX over many cellular responses including cell migration and communication, previously only attributed to gap junction blockade, are indeed the consequence of the disruption of intracellular Ca²⁺ homeostasis.

**One Sentence Summary:** carbenoxolone blocks cell migration by inhibiting SERCA

## INTRODUCTION

Collective and single cell migration are fundamental for tissue morphogenesis and repair, immunosurveillance, angiogenesis, and cancer progression (Abu Khamidakh et al., 2013; Cao, 2025; Friedl & Gilmour, 2009; Scarpa & Mayor, 2016). Cellular locomotion is a dynamic process that requires the asymmetric distribution of intracellular components, including the organelles, in a process termed cellular polarization. Consequently, changes in the location and/or the function of the organelles directly affects motility (Marwaha & Das, 2026). During collective cell migration leader cells at the migratory front transfer signals using extra and intracellular communication mechanisms to ensure coordinated behavior across the migrating monolayer. This includes the transference of Ca^2+^ waves, which at cellular level control several components of the migration machinery, such as cell adhesion, actomyosin contractility, tension, and membrane dynamics (Brun-Cosme-Bruny et al., 2026; Tsai et al., 2015). However, how cells coordinate intracellular signaling and intercellular communication remains a central question.

Organelle Ca^2+^ signaling plays a pivotal role in several processes including cell migration (Tsai et al., 2015). The ER serves as the primary reservoir of intracellular Ca^2+^, capable of accumulating high concentrations, reaching millimolar levels (Barrero et al., 1997). This is achieved through the interaction of three key components: 1) Ca^2+^ ATPases from the sarco(endo)plasmic reticulum Ca^2+^ ATPase (SERCA) family, localized in the ER membrane, which actively transport Ca^2+^ against its concentration gradient; 2) Ca^2+^-binding proteins present in the ER lumen, such as calreticulin and calsequestrin, which buffer the stored Ca^2+^; and 3) Ca^2+^ channels, such as inositol 1,4,5-trisphosphate receptors (IP_3_Rs) and the ryanodine receptors (RyRs), that release stored Ca^2+^ into the cytosol along its concentration gradient. Consequently, intercellular Ca²⁺ waves initiated by mechanical or chemical stimuli are mainly derived from the ER (Leybaert & Sanderson, 2012), and proper maintenance of intraluminal Ca^2+^ levels is critical for maintaining ER functions. Prolonged depletion of Ca^2+^ in the ER triggers the unfolded protein response (UPR), leading to ER stress that distorts cell migration (Sáez et al., 2014), and could result in apoptosis (Carreras-Sureda et al., 2018). Therefore, SERCA inhibitors such as thapsigargin and di-tert-butylhydroquinone (BHQ) are widely used to induce UPR and are considered potential therapeutic agents in cancer treatment (Mahalingam et al., 2016). Consequently, SERCA function and intercellular communication are tightly interconnected, to provide efficient signal transduction and cellular coordination (Ho et al., 2021).

Several cellular communication mechanisms contribute to ensure effective coordination during collective cell migration. Connexin (Cxs) and Pannexin (Panxs) forming channels, often named hemichannels (HCs), and Panx channels, allow autocrine and paracrine communication, respectively. At the plasma membrane these proteins form membrane channels that allow uptake and release of molecules from the extracellular milieu, or dock to similar channels from an adjacent cell to form gap junction channels (GJCs), which allow direct intracellular communication (Martins-Marques et al., 2025; Palacios-Prado et al., 2022; Peters et al., 2007; Saez et al., 1989). GJCs allow efficient transference of ions (i.e. Ca^2+^), second messengers, small metabolites, and small peptides and RNAs (Harris, 2007). Consequently, GJCs are associated with various functions such as ischemic brain damage, immunity, hippocampal plasticity or propagation of Ca^2+^ waves in several cell types. The relative contribution of each of Cxs and Panxs in these responses has been frequently assessed by using pharmacological inhibitors (Willebrords et al., 2017). However, the individual contribution of these channels during collective cell migration remains unclear due to the broad effect of most of these compounds (Abu Khamidakh et al., 2013; Cao, 2025; Friedl & Gilmour, 2009; Harcha et al., 2021; Song et al., 2023; Walker et al., 1994).

Carbenoxolone (3β-hydroxy-11-oxoolean-12-en-30-oic acid 3-hemisuccinate, CBX), a water-soluble synthetic derivative of 18β-glycyrrhetinic acid (18β-GA) extracted from licorice root (*Glycyrrhiza glabra*), has been extensively used as a broad-spectrum pharmacological tool to inhibit Cx- and Panx-mediated intercellular communication. CBX reduces cell migration in several cellular models, such as glioblastoma (Oliveira et al., 2005), breast cancer (Pollmann et al., 2005), and others (Harcha et al., 2021). Whereas these studies suggest the contribution of GJC-mediated communication, the conclusions are exclusively based on the use of this drug that also blocks Panx-forming channels (Michalski & Kawate, 2016). Importantly, CBX, and other glycyrrhetinic acids, are used to study the role of Cx and Panx channels, but pharmacological specificity of CBX has been increasingly questioned, and off-target effects have been reported (Table 1). For example, the direct and indirect inhibition of 11β-hydroxysteroid dehydrogenase by CBX has been proposed (Devang et al., 2023), but not fully demonstrated. Moreover, CBX blocked Ca²⁺ waves in a time-dependent manner by inhibiting IP_3_-evoked Ca²⁺ release rather than by blocking GJCs per se, suggesting a significant off-target action on endoplasmic reticulum (ER) Ca^2+^ signalling (Buckley et al., 2021). Furthermore, CBX and 18β-GA were shown to depolarize the mitochondrial membrane potential, suggesting that their effects on cell behavior may be mediated, at least in part, through disruption of intracellular Ca^2+^ homeostasis independently of gap junction blockade. These findings suggest that the effects of CBX in different multicellular processes, such as collective cell migration, are a combined effect of blocking intercellular communication and altering organelle-level Ca^2+^ signalling.

**Table 1.**
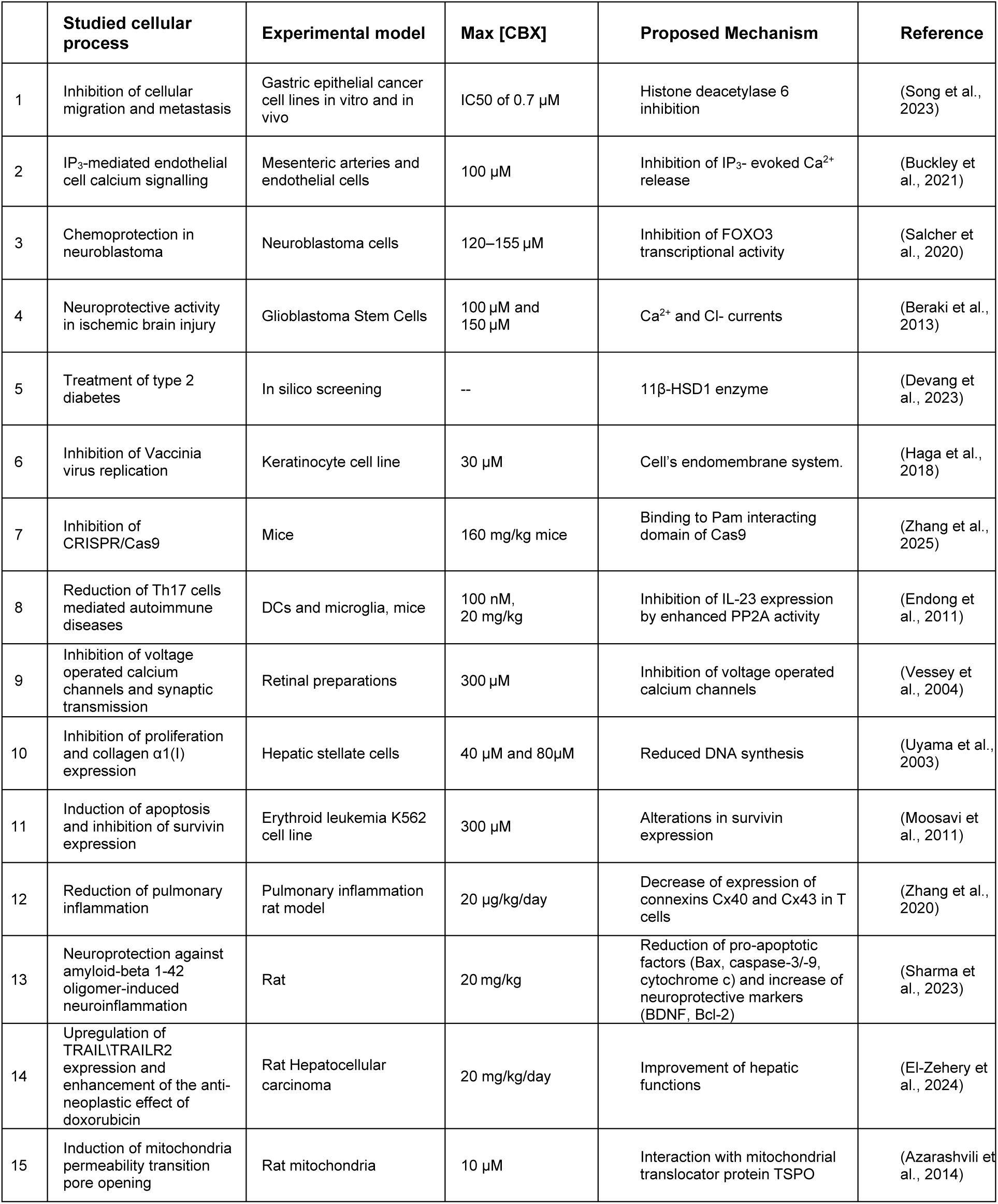
Reported CBX effects independent of inhibition of gap junctions.

|  | Studied cellular process | Experimental model | Max [CBX] | Proposed Mechanism | Reference |
| --- | --- | --- | --- | --- | --- |
| 1 | Inhibition of cellular migration and metastasis | Gastric epithelial cancer cell lines in vitro and in vivo | IC50 of 0.7 μM | Histone deacetylase 6 inhibition | (Song et al., 2023) |
| 2 | IP<sub>3</sub>-mediated endothelial cell calcium signalling | Mesenteric arteries and endothelial cells | 100 μM | Inhibition of IP<sub>3</sub>- evoked Ca<sup>2+</sup> release | (Buckley et al., 2021) |
| 3 | Chemoprotection in neuroblastoma | Neuroblastoma cells | 120–155 μM | Inhibition of FOXO3 transcriptional activity | (Salcher et al., 2020) |
| 4 | Neuroprotective activity in ischemic brain injury | Glioblastoma Stem Cells | 100 μM and 150 μM | Ca<sup>2+</sup> and Cl- currents | (Beraki et al., 2013) |
| 5 | Treatment of type 2 diabetes | In silico screening | -- | 11β-HSD1 enzyme | (Devang et al., 2023) |
| 6 | Inhibition of Vaccinia virus replication | Keratinocyte cell line | 30 μM | Cell's endomembrane system. | (Haga et al., 2018) |
| 7 | Inhibition of CRISPR/Cas9 | Mice | 160 mg/kg mice | Binding to Pam interacting domain of Cas9 | (Zhang et al., 2025) |
| 8 | Reduction of Th17 cells mediated autoimmune diseases | DCs and microglia, mice | 100 nM, 20 mg/kg | Inhibition of IL-23 expression by enhanced PP2A activity | (Endong et al., 2011) |
| 9 | Inhibition of voltage operated calcium channels and synaptic transmission | Retinal preparations | 300 μM | Inhibition of voltage operated calcium channels | (Vessey et al., 2004) |
| 10 | Inhibition of proliferation and collagen α1(I) expression | Hepatic stellate cells | 40 μM and 80μM | Reduced DNA synthesis | (Uyama et al., 2003) |
| 11 | Induction of apoptosis and inhibition of survivin expression | Erythroid leukemia K562 cell line | 300 μM | Alterations in survivin expression | (Moosavi et al., 2011) |
| 12 | Reduction of pulmonary inflammation | Pulmonary inflammation rat model | 20 μg/kg/day | Decrease of expression of connexins Cx40 and Cx43 in T cells | (Zhang et al., 2020) |
| 13 | Neuroprotection against amyloid-beta 1-42 oligomer-induced neuroinflammation | Rat | 20 mg/kg | Reduction of pro-apoptotic factors (Bax, caspase-3/-9, cytochrome c) and increase of neuroprotective markers (BDNF, Bcl-2) | (Sharma et al., 2023) |
| 14 | Upregulation of TRAIL/TRAILR2 expression and enhancement of the anti-neoplastic effect of doxorubicin | Rat Hepatocellular carcinoma | 20 mg/kg/day | Improvement of hepatic functions | (El-Zehery et al., 2024) |
| 15 | Induction of mitochondria permeability transition pore opening | Rat mitochondria | 10 μM | Interaction with mitochondrial translocator protein TSPO | (Azarashvili et al., 2014) |

In the present study, we show that CBX reduces cell migration as a consequence of the fast and reversible inhibition of the SERCA pump that leads to a passive leak of the ER Ca^2+^ store. We propose that some of the effects of CBX described in the literature, previously attributed to Cx- and Panx-forming channels, are instead a consequence of the impact on ER Ca^2+^.

## RESULTS

### Carbenoxolone reduces cell migration by disrupting ER Ca^2+^

Cellular coordination and signaling during collective migration have been described at different levels (Kotini & Mayor, 2015; Sharma et al., 2023; Vitorino & Meyer, 2008), but whether there is an interplay between cellular communication and organelle signaling during this process remains unknown. We used Human Umbilical Vein Endothelial Cells (HUVECs) as a model of collective cell migration and found that broad blockade of Cx and Panx channels (i.e. GJCs and HCs) with CBX reduced the speed of migration in a dose-dependent manner (Fig. 1A and B, and Fig. Supplement 1A–C). By contrast, the blockade of Cx HCs with La^3+^, and Panx channels with probenecid had no effect (Fig. 1B), suggesting that GJCs but not HCs contribute to collective cell migration. Interestingly, the reduction observed with 50 µM CBX phenocopied the results obtained with thapsigargin (Fig. 1A and B and Fig. Supplement 1A). This effect was observed very early upon treatment (Fig. Supplement 1C) and was also observed during collective migration of ARPE-19 epithelial cells (Fig. Supplement 1D).

**Fig. 1.**
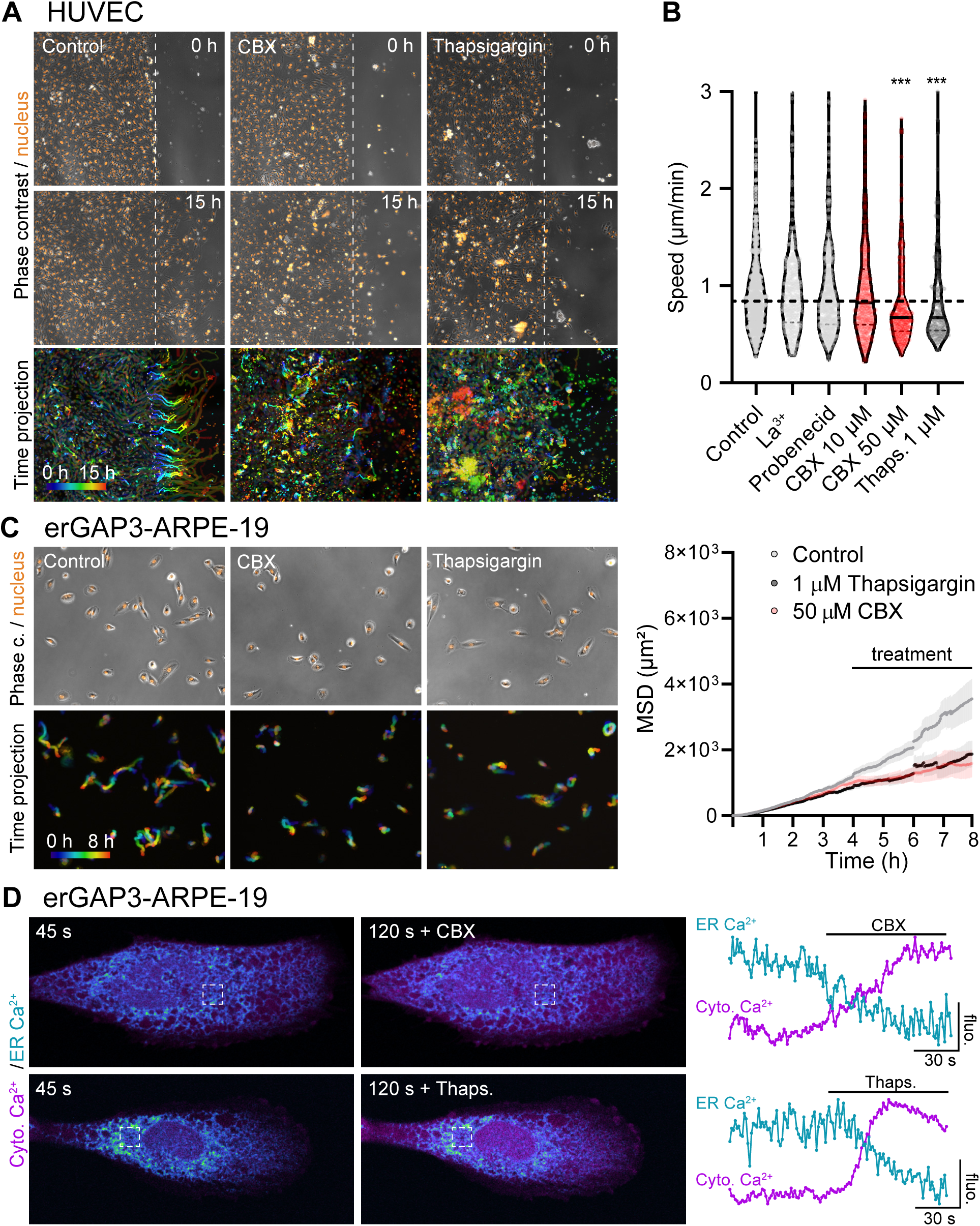
Collective cell migration depends on ER Ca^2+^. **(A)** Microscopy images showing nuclear staining (Hoechst; orange) and phase contrast during collective migration of HUVEC cells. The scratch is denoted by the dotted line at indicated times. The bottom row shows a time-projection image in which Control cells, but not cells treated with carbenoxolone (CBX; 50 µM) and thapsigargin (1 µM), show directional tracks. (**B)** Graph showing the instantaneous speed of migration in one of four independent experiments (n=4; n>250 cells per condition, per experiment). The line denotes the median. ***p<0.01 Kruskal-Wallis test compared to Control condition. **(C)** Microscopy images of a 2D single cell migration of erGAP3-ARPE-19 cells, under resting conditions or treated with CBX or thapsigargin. Time projection of 8 h of migration. The graph shows the mean square displacement (MSD) under resting conditions or after acute treatment with CBX or thapsigargin. **(D)** Simultaneous imaging of cytosolic (Calbryte 590) and ER (erGAP3) Ca^2+^ by super-resolution microscopy upon addition of thapsigargin (1 µM) or CBX (50 μM) in erGAP3-ARPE-19 cells. Representative images at indicated times 45s before and 120s after the treatment (+ Thaps., or + CBX) are shown.

HUVEC collective cell migration was quickly reduced by CBX and thapsigargin, suggesting that this effect is due to changes in intercellular communication and/or signaling, but unlikely ER stress or similar ER dysfunctions with slower kinetics. To support this, we tested the acute effect over migration using 2D single cell migration, in which there are no or few transient cellular contacts (i.e. no GJCs). We found that upon treatment both CBX and thapsigargin, quickly reduced the migratory behavior of ARPE-19 epithelial cells (Fig. 1C). Since in these conditions there are few or no GJCs, we concluded that CBX might have an effect independent of GJCs, and that it may have the same molecular target as thapsigargin.

### Carbenoxolone evokes Ca^2+^ release from the ER

Since thapsigargin empties the ER Ca^2+^ store which leads to an increase in cytosolic Ca^2+^, we hypothesized that the effect of CBX over cell migration could be related to changes in ER and cytosolic Ca^2+^. Simultaneous monitoring of ER and cytosolic Ca^2+^ using super-resolution microscopy showed that CBX provoked a decrease in the ER Ca^2+^ and a concomitant increase in the cytosolic Ca^2+^, and these effects were very similar to those provoked by thapsigargin (Fig. 1D). Altogether, these data suggest that the molecular targets of CBX and thapsigargin are related.

In order to gain more insight into the mechanism of action of CBX, we monitored ER and cytosolic Ca^2+^ with the ratiometric Ca^2+^ indicators erGAP3 and fura-2, respectively. CBX evoked a decrease of the ER Ca^2+^ that was accompanied by a cytosolic Ca^2+^ transient, and these effects were reversible upon CBX washout (Fig. 2A). Comparison of Ca^2+^ kinetics in the two compartments showed that the cytosolic Ca^2+^ peak preceded the minimum ER Ca^2+^ decrease, as previously described in various cell types (Navas-Navarro et al., 2016). The removal of CBX provoked a small transient of cytosolic Ca^2+^, probably due to store-operated Ca^2+^ entry (SOCE) activation. We found that the effect of CBX on the ER Ca^2+^ was common to several cell types, such as HeLa (Fig. 2A), retinal epithelial ARPE-19 (Fig. 2B), HEK293 cells (Fig. 2C), and primary mouse astrocytes isolated from transgenic mice expressing erGAP3 (Fig. 2D). The effect of CBX on the ER-Ca^2+^ was concentration-dependent between 10 and 100 μM (Fig. 2C). Moreover, the ER Ca^2+^ depletion was slower than that evoked by inositol-trisphosphate (IP_3_)-producing agonists such as ATP together with carbachol (CCh) (Fig. Supplement 2). These results indicate that the effect of CBX does not involve membrane receptor activation and/or IP_3_R cascade amplification and that CBX passively empties the ER Ca^2+^ store.

**Fig. 2.**
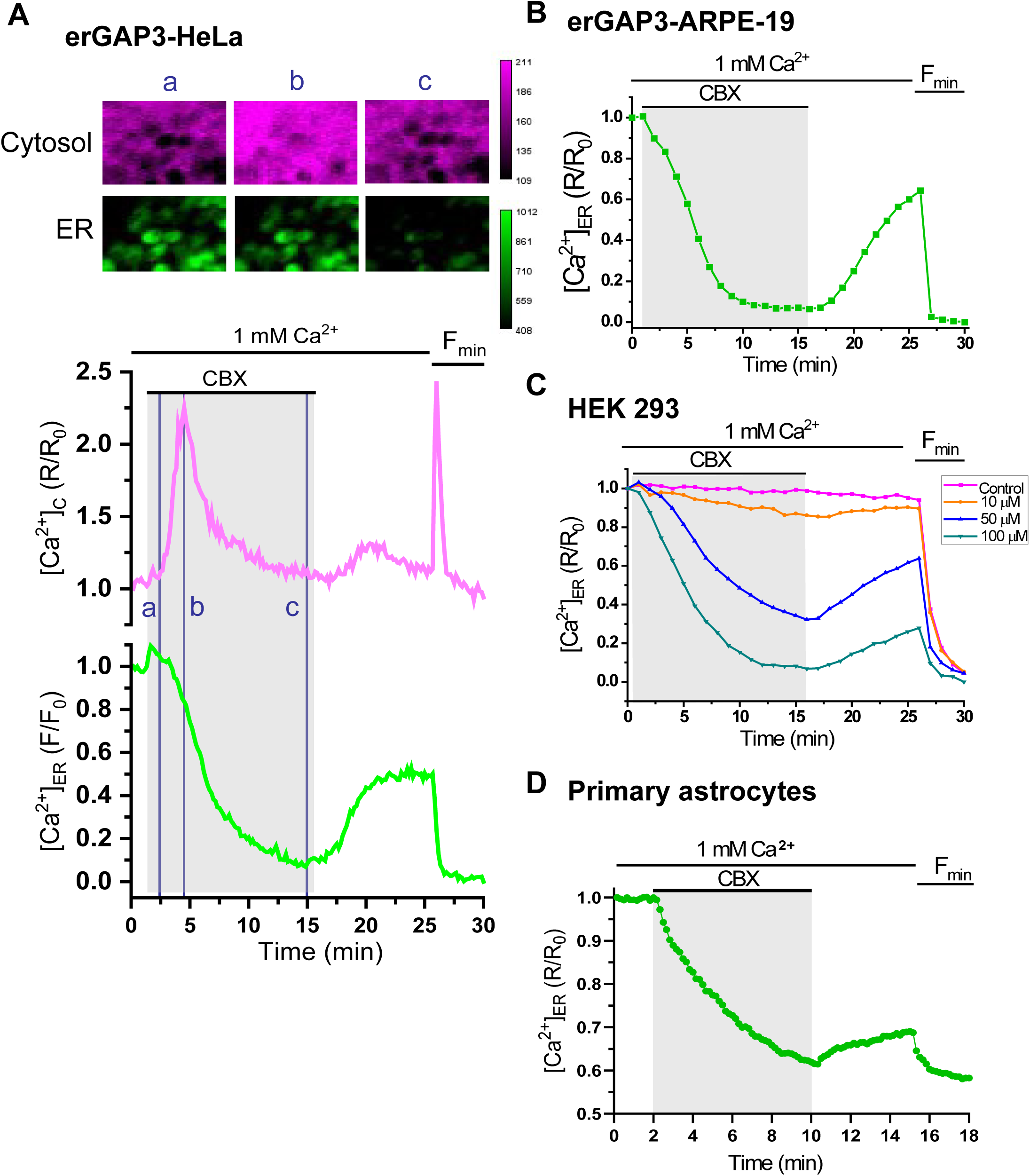
CBX empties the ER of Ca^2+^ in various cell types. Depletion of ER Ca^2+^ by treatment with CBX (100 μM) in erGAP3-HeLa **(A)**, erGAP3-ARPE-19 **(B),** HEK293 cells **(C)**, or mouse cortical astrocytes **(D)**, all expressing the ER Ca^2+^ indicator erGAP3. **(A)** Simultaneous imaging of cytosolic (fura-2; above) and ER (erGAP3; below) Ca^2+^ responses upon CBX (100 μM) addition to HeLa cells. Fura-2 is expressed as F340/F380 ratio (R) normalized (R/R_0_). Each trace is the mean of all cells (91) in the optical field. The three pseudocolor images (a,b,c) are captured at the times indicated in the graph. Calibration of GAP3 was obtained by adding a depletion cocktail (indicated by Fmin) containing ATP (100 µM), the membrane-permeable SERCA inhibitor 2,5-di-tert-butylhydroquinone (BHQ, 10 μM) and EGTA (0.5 mM). **(B)** Traces shown are the average values of 42 cells from the same microscope field and are representative of 4 similar experiments. [Ca^2+^]_ER_ is represented as F470/F405 ratio (R) normalized to R_0_ (R/R_0_). GAP3 is expressed as F470 normalized to F/F_0_. **(C)** Dose-response to three CBX concentrations (10, 50 and 100 μM) on the ER Ca^2+^ release in HEK293 cells. Traces are the average of 130 cells (Control), 164 cells (10 μM), 122 cells (50 μM) and 95 cells (100 μM). **(D)** Effect of CBX (100 μM) on the ER Ca^2+^ release from the ER of mouse cortical astrocytes isolated from erGAP3 transgenic mice. The trace is the mean of 14 cells.

### Carbenoxolone inhibits SERCA activity

To gain more direct insight into the molecular mechanisms underlying the effect of CBX on ER Ca^2+^, we used a model that provides direct access to the ER consisting of digitonin-permeabilized cells (Fig. 3A). Thus, we used cells expressing a luminescent low affinity Ca^2+^ indicator targeted to the ER lumen (IgGAP1) to quantify ER Ca^2+^ (Rodriguez-Prados et al., 2015). We used two protocols to directly evaluate the effect of CBX on SERCA activity. In the first approach, the ER was emptied of Ca^2+^, and then Ca^2+^ (200 nM) was perfused in the absence (Control) or presence (CBX) of CBX. The ER Ca^2+^ uptake by SERCA pump was practically abolished by CBX (Fig. 3B and 3C). A second protocol to assess the CBX effect was conducted when the ER Ca^2+^ store was full. The treatment with CBX (100 μM) provoked a strong and progressive reduction of the ER Ca^2+^ that practically emptied the ER Ca^2+^ store (Fig. 3D). The CBX-induced ER Ca^2+^ depletion was reversible and had an EC_50_ of 66.5 µM (Fig. 3E). In agreement with results in intact cells, the effect of CBX was similar to that evoked by the specific reversible inhibitor of SERCA BHQ (10 μM) (Fig. 3F and 3G). Moreover, subsequent addition of BHQ during the slow exit of Ca^2+^ from the ER evoked by CBX had no additional effect, indicating that both agents are acting on the same molecular target (Fig. 3H and Supplement 3). Similar findings were obtained with CBX precursor glycyrrhetinic acid (GA), which also provoked a Ca^2+^ release from the ER with similar kinetics to that induced by CBX (Fig. Supplement 4).

**Fig. 3.**
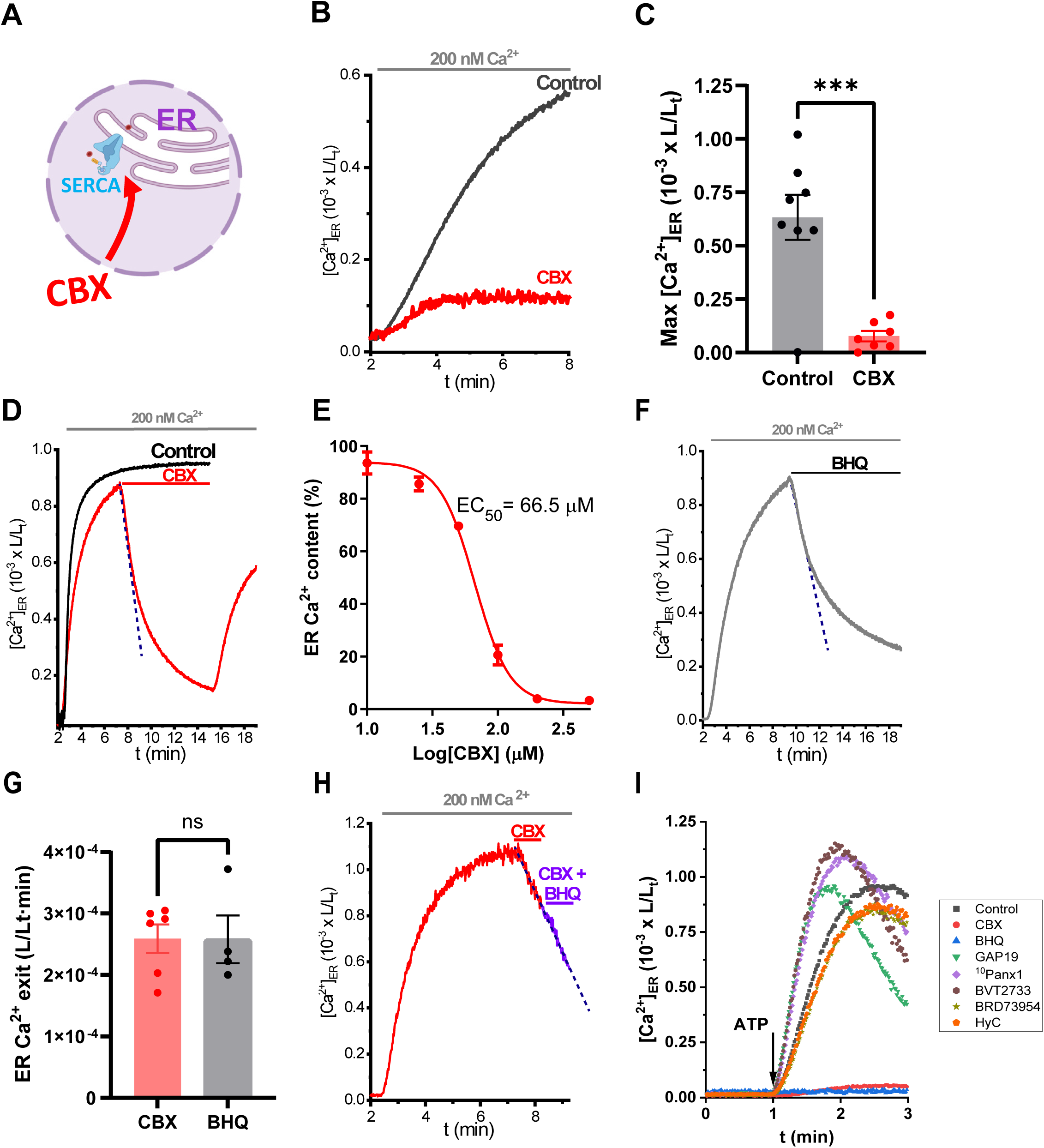
Direct monitoring of ER Ca^2+^ depletion by CBX in permeabilized cells. **(A)** Schematic drawing of experimental design. **(B)** Active Ca^2+^ uptake by the ER in digitonin-permeabilized HeLa cells expressing er(Ig)GAP1. Cells were perfused with an intracellular-like buffer containing 200 nM Ca^2+^, in the absence (Control), or the presence of CBX (100 µM). Each trace is the average of 4×10^4^ cells. **(C)** Pooled data of similar experiments to that shown in **B**. Each point represents the maximal [Ca^2+^]_ER_ obtained at min 8 upon Ca^2+^ addition. Data are expressed as mean ± SEM. A Student′s *t* test was applied. ***p <0.001. **(D)** er(Ig)GAP1 expressing HeLa cells were permeabilized and perfused at minute 8 with CBX (100 µM; red trace) until minute 15 when CBX was washed out. (**E)** Concentration-dependent response on the ER Ca^2+^ content (expressed as percentage of the total content of Ca^2+^) upon CBX addition. Data are represented as mean ± SEM (n = 3–4). (**F)** Passive ER Ca^2+^ depletion with BHQ (10 μM). Experimental conditions were identical to D. (**G)** Pooled data quantifying the ER Ca^2+^ exit, monitored as L/Lt·min (mean ± SEM from 4-6 experiments). A Student′s *t* test was applied. (**H)** Comparison of the release evoked by CBX (100 µM) alone or together with BHQ (10 µM) in HEK293 cells expressing erGAP3. Traces represent the average luminescent signal from 1–1.5 × 10⁵ cells. **(I)** Comparison of the effect of CBX with other inhibitors that act on alternative targets of CBX on SERCA activity. Stably expressing er(Ig)GAP1-HeLa cells were permeabilized in an intracellular-like Ca^2+^-free medium with digitonin and perfused with an intracellular-like medium containing 200 nM Ca^2+^ without ATP. Then, inhibitor was added and incubated for 10 min prior to injecting ATP (1 mM) to trigger Ca^2+^ uptake into the ER. At the end of each run, a second injection with saturating Ca^2+^ (10 mM) was added to calibrate the luminescent signal. The inhibitors used were: GAP19, ^10^Panx1, hydrocortisone cypionate (HC), BVT2733, BRD73954, CBX (all at 100 µM), and BHQ (10 µM). Each point is the average of 3-10 wells, each with 7 x 10^3^ cells.

It has been shown that CBX inhibited IP_3_-evoked Ca^2+^ signalling (Buckley et al., 2021). Consequently, we evaluated whether CBX inhibits IP_3_R-mediated Ca^2+^ responses. We used cells lacking IP_3_R-mediated signaling (triple IP_3_R KO HEK293 cells) (Alzayady et al., 2016), to fully rule out the contribution of IP_3_Rs. We observed that CBX emptied the ER of Ca^2+^ with similar kinetics in both wild type and IP_3_R KO cells (Supplement Fig. 5). These results indicate that the CBX-evoked Ca^2+^ exit does not involve IP_3_Rs.

CBX is best known as a Cx- and Panx-channel blocker, but other potential targets have also been reported (Table 1), such as 11β-hydroxy-steroid dehydrogenase type 1, and histone deacetylase 6. Because Cx and Panx channels are permeable to Ca^2+^ (Harris, 2007), we evaluated the effects of several well-established inhibitors acting on alternative targets of CBX on ER Ca^2+^ in permeabilized cells (Fig. 3I). We used BVT2733 and hydrocortisone cypionate (HyC) that inhibit 11β-hydroxy-steroid dehydrogenase type 1 (11β-HSD1); BRD73954, that inhibits histone deacetylase 6 (HDAC6); and, ^10^Panx1 and GAP19, which inhibit Panx1 and Cx43 channels, respectively (Supplement Fig. 6). Importantly, we found that none of these agents modified SERCA activity, which was fully inhibited by CBX and BHQ (Fig. 3I).

Altogether, these data demonstrate that the effect of CBX on ER Ca^2+^ is produced by directly inhibiting SERCA activity, and not secondary to the potential effect of CBX on alternative targets.

### Carbenoxolone directly binds SERCA

We next investigated the intracellular localization of CBX to gain further evidence of its function. First, we synthesized a fluorescent derivative CBX by fusing it with a BODIPY molecule which we named CBX-A (Supplement Fig. 7). We labelled HUVEC cells with CBX-A and co-stained them with several markers. Immunofluorescence analysis showed that CBX-A partially co-localized with Panx1 at the plasma membrane (Fig. 4A-4C), as previously described (Michalski & Kawate, 2016), but also co-localized with an ER marker (Nogo/Rtn4), suggesting its binding to an ER resident protein like SERCA. We indeed confirmed that CBX-A co-localized with SERCA2b, both in intact and in permeabilized HEK293 cells (Fig. 4D and E).

**Fig. 4.**
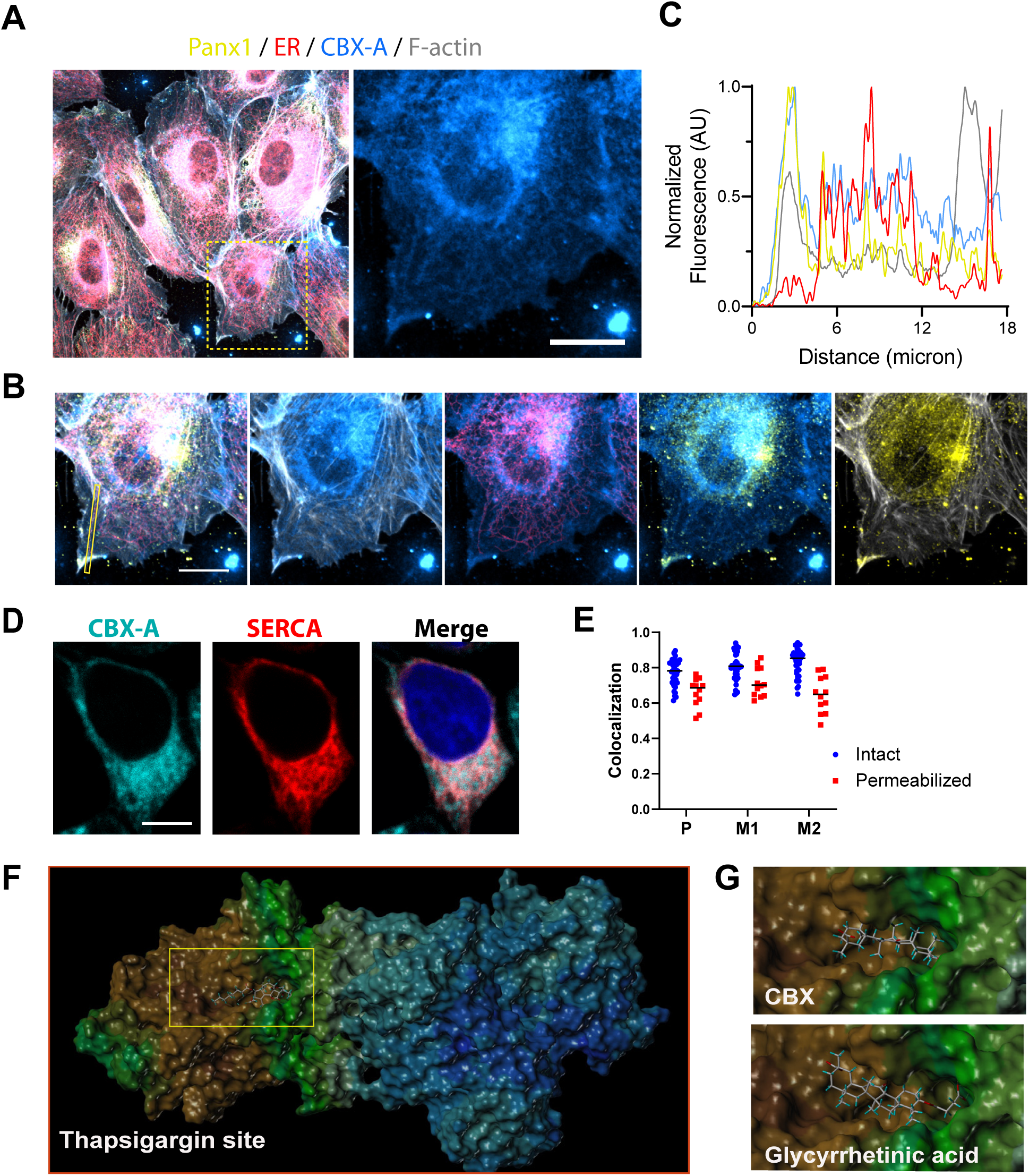
CBX binds to SERCA. **(A)** Microscopic images of HUVEC cells stained with CBX-A (50 µM; cyan). Panx1 plasma membrane channel (yellow), ER resident protein NoGo-A/Rtn4-A (red), and F-actin (white) labeling is also shown. (**B)** Zoom picture of the inset shown in **A**. From left to right: CBX-A / ER / Panx1; CBX-A / F-actin; CBX-A / ER; CBX-A / Panx1; and Panx1 / F-actin. (**C)** Quantification of colocalization within the line scan indicated in yellow in the first panel in **B**. Note that CBX-A co-localizes with Panx1 at the plasma membrane as well as with the ER. (**D)** Colocalization of CBX-A with SERCA2b in HEK293 cells. Representative confocal images of live HEK293 expressing cherry-SERCA2b (red) incubated with CBX-A (50 µM; cyan) for 15 min. (**E)** Quantitative colocalization analysis of confocal fluorescence microscopy images of either live or fixed cells, expressed as Pearson’s (P) coefficient, Mandeŕs coefficient 1 (M1; Serca/CBX) and Mandeŕs coefficient 2 (M2; CBX/Serca). Each point indicates one cell analysed from 3 independent experiments. (**F)** Molecular docking analysis of SERCA. The complex is shown with a surface display encoding the lipophilic potential (brown, lipophilic; blue, hydrophilic). The transmembrane part of the complex stands out with its lipophilic region shown in brown. The binding site of the inhibitor thapsigargin (shown as stick model) is indicated (yellow square). (**G)** Molecular docking modelling for binding of CBX or glycyrrhetinic acid to SERCA1.

To gain further insight into the putative binding site of CBX within the SERCA molecule, we performed molecular docking analysis. Our results revealed that CBX exhibited a favorable affinity toward SERCA, occupying binding pocket of the canonical SERCA inhibitor thapsigargin within the transmembrane domain (Fig. 4F and G). Docking predictions of the CBX-SERCA1 complex showed that CBX, and also GA adopted a well-defined orientation within the binding cavity of thapsigargin.

We conclude that the depletion effect of CBX on ER Ca^2+^ is due to its direct binding to the SERCA protein localized in the ER.

## DISCUSSION

Cell-to-cell communication mediated by Cx- and Panx-forming channels is crucial to coordinate and synchronize physiological and physiopathological responses in many tissues and organs (Harris, 2007). To study these pathways, carbenoxolone (CBX) and its precursor molecule, glycyrrhetinic acid, have long been instrumental as widely used standard blockers (Koval et al., 2023). However, our findings demonstrate that CBX exerts a profound, channel-independent effect on intracellular calcium homeostasis. By utilizing the ratiometric erGAP Ca^2+^ sensor family for precise quantification, we show that CBX provokes a rapid depletion of the endoplasmic reticulum (ER) Ca^2+^ store, accompanied by a concomitant increase in cytosolic Ca^2+^. This phenomenon was not cell-type specific; it was consistently observed across multiple cell types, indicating that the underlying molecular target of CBX leading to these ER alterations is ubiquitously expressed.

The steady-state of the ER Ca^2+^ store relies on a strict balance between active uptake by the sarco-endoplasmic reticulum Ca^2+^-ATPase (SERCA) pump—a P-type ATPase superfamily member transporting two Ca^2+^ ions per ATP hydrolyzed—and passive efflux through leak channels (Burdakov et al., 2005). The kinetics of the ER Ca^2+^ depletion induced by CBX closely mirror the passive emptying triggered by established SERCA inhibitors like thapsigargin, BHQ, or cyclopiazonic acid (Navas-Navarro et al., 2016; Rodriguez-Prados et al., 2015), while differing drastically from the transient Ca^2+^ release triggered by IP_3_-coupled agonists.

Our evidence supports a mechanism of direct physical interaction and inhibition. While labeled CBX binds to targets at the plasma membrane—consistent with its known blockade of Cxs and Panxs (Matuseviciute et al., 2023; Michalski & Kawate, 2016; Willebrords et al., 2017)— was also detected in the ER, where it directly co-localizes with SERCA. This is corroborated by our molecular docking data, predicting that both CBX and glycyrrhetinic acid fit into the well-defined transmembrane pocket utilized by thapsigargin in its E2 state (Obara et al., 2005), thereby blocking Ca^2+^ transport across the ER membrane. Functionally, two independent SERCA activity assays (triggered by Ca^2+^ or extracellular ATP) confirmed that CBX completely abolishes Ca^2+^ transport into the ER lumen, phenocopying BHQ. Crucially, while a growing number of alternative enzymatic targets for CBX have been proposed (such as 11β-HSD1 or HDAC6A, see Table 1), using specific inhibitors against these enzymes failed to affect SERCA activity, ruling out their involvement and confirming a direct inhibitory mechanism on the pump. Collective cell migration relies on the precise coordination of cells moving across a monolayer, which depends on diverse cellular communication mechanisms (Harcha et al., 2021; Hino et al., 2020). Cxs and Panxs promote extracellular and intracellular cell communication to assist migration (Harcha et al., 2021; Pun et al., 2024; Sharma et al., 2023), but channel-independent effects on motility have also been reported (Koepple et al., 2021). CBX is frequently used to monitor the role of GJCs and HCs on motility (Harcha et al., 2021; Liu et al., 2019; Oliveira et al., 2005; Pollmann et al., 2005), but the effects over motility has been disputed as indirect evidence points to distinct alternative targets (Karsch-Bluman et al., 2017; Salcher et al., 2020; Song et al., 2023).

We show that CBX reduces collective cell migration by leaking Ca^2+^ from the ER, which cannot be refilled, likely preventing intercellular Ca^2+^ wave transference. Proper SERCA function regulates ER Ca^2+^ content, cytosolic Ca^2+^ levels, and plasma membrane permeability via SOCE. Chronic alterations in ER Ca^2+^ levels impair protein synthesis, folding, and secretion, triggering ER stress and eventually apoptosis (Mekahli et al., 2011). Induction of ER stress reduces cell migration in most healthy cells, including endothelial cells (Sáez et al., 2014). This pathway, alongside disturbed ER and cytosolic Ca^2+^, could lower actomyosin contractility to prevent motility (Sáez et al., 2018). However, because CBX rapidly impairs collective migration, this acute motility defect likely relies on disturbed Ca^2+^ signaling rather than downstream ER stress, which develops with slower kinetics. To further test whether this acute impairment depends on cell-cell communication, we evaluated 2D single-cell migration in conditions with few or no transient cellular contacts (and thus no functional GJCs). The rapid reduction in migratory behavior observed with both CBX and thapsigargin under these conditions confirms that CBX impairs migration through a mechanism independent of GJC blockade, acting on the same molecular pathway as SERCA pump inhibition.

These findings require a careful reevaluation of previous literature where reductions in cell motility, or network function caused by CBX were attributed solely to the blockade of GJCs or HCs. Electrophysiological, pharmacological, and cryo-EM studies conclusively demonstrate that CBX binds and blocks Panx1 channels (Ruan et al., 2020) and Cxs (Matuseviciute et al., 2023; Michalski & Kawate, 2016; Willebrords et al., 2017). Nonetheless, indirect evidence previously hinted at alternative targets: CBX inhibits spontaneous network activity in astrocytes even in cultures derived from *Cx43* knockout mice (Rouach et al., 2003), and its anti-epileptic activity appears unrelated to GJCs (Beaumont & Maccaferri, 2011).

Consequently, conclusions obtained in experimental setups relying exclusively on broad and unspecific blockers can be highly misleading. While communication-related proteins like Cx43 and Panx1 actively coordinate Ca^2+^ and ERK signaling (Koepple et al., 2021; Liu et al., 2019; Sáez et al., 2014), their specific roles must be verified using multiple blockers alongside genetic tools (Harris, 2007; Sáez et al., 2017). To advance the field, future studies must routinely temper and contrast the contribution of these pathways against the dual effects of CBX, carefully distinguishing cell communication blockade from direct SERCA pump inhibition.

## MATERIALS AND METHODS

### Cells

The two stably-expressing erGAP3-HeLa and erGAP3-ARPE-19 cell lines were used for Ca^2+^ imaging experiments to monitor ER Ca^2+^ and have been previously described (Navas-Navarro et al., 2016; Rojo-Ruiz et al., 2024). Wild-type HeLa, HEK-293T and HEK-293 cells engineered using CRISPR-Cas9 technology to knock out the three IP_3_ receptors (3IP_3_R KO) were also used (Alzayady et al., 2016). The stably expressing IgGAP1-Hela cell lines was used to perform the ER Ca^2+^ measurements in batch by luminescence and have been described earlier (Rodriguez-Prados et al., 2015). HeLa cells and HEK293T cells were maintained in Dulbecco’s modified Eagle’s medium (DMEM) and ARPE-19 cells were maintained in DMEM/F12 (1:1), both supplemented with 10% (v/v) heat inactivated fetal bovine serum, GlutaMAX supplement and 100 μg/ml streptomycin and 100 U/ml penicillin (all Invitrogen). The stable clones were maintained in the presence of 200 µg/ml G418 (Gibco). All cells were maintained at 37°C in a humidified atmosphere with 5% CO_2_ and were passaged by trypsinization. For single cell imaging, cells were seeded onto 12 mm-coverslips coated with poly-L-lysine (Sigma; 0.1% (w/v)). Human umbilical vein endothelial cells (HUVECs, PromoCell, C-12203) were cultured in ready-to-use full endothelial growth medium (PromoCell C-22010), that included the Basal Medium and SupplementMix, and was supplemented with 1% Penicillin-Streptomycin (Pen-Strep, ThermoFisher, P06-07100) and plated on Corning culture dishes with 100 mm diameter (Sigma-Aldrich, CLS430167) coated with fibronectin from bovine plasma (1 µL/mL, Sigma-Aldrich, F1141). HUVECs were maintained at 37°C and 5% CO_2_ for 2 days until a ∼90% confluency was reached, in preparation for the next passage.

Astrocytes were isolated from postnatal day P2-3 C57BL/6JCrl (Charles Rivers) erGAP3 (L10) transgenic mouse pups (Navas-Navarro et al., 2016). The cortices were dissected in Hank’s balanced salt solution (HBSS; Gibco™, ThermoFisher,) supplemented with 10 mM glucose and 0.05% (w/v) BSA, cut into small pieces and digested for 20 min at 37°C with 0.5 mg/ml papain (Worthington Biochemical Corporation) and 0.04 mg/ml DNase (Roche). Cells were resuspended in Neurobasal™ medium, supplemented with 1% glutaMAX™-I (ThermoFisher, Gibco™); 2 % (v/v) B27™ (ThermoFisher, Gibco™); 10.000 U/mL penicillin-streptomycin; and 10% (v/v) horse serum (HS; ThermoFisher, Gibco™); and plated on a poly-L-lysine-coated 75 cm^2^ flask. At days 2 and 5, flasks were slapped to remove neurons and other loosely attached cells such as microglia and oligodendrocytes. After 1 week, cells were trypsinized and seeded in the same medium onto poly-L-lysine-coated 12 mm diameter glass coverslips at a density of 2.5 x 10^4^ cells/coverslip. Cultures were used for imaging experiments within 1–2 weeks.

### Plasmids

pcDNA3-erGAP3 or pcDNA3-er(Ig)GAP1 plasmids were transiently transfected in wild-type HEK293, HeLa or 3IP_3_KO HEK293 cells, using Lipofectamine® 2000 following manufacturer’s instructions and used for imaging experiments 24-48 hrs after transfection. pcDNA3-Cherry-Serca2b has been previously reported (Manjarres et al., 2010).

### Reagents

Carbenoxolone disodium salt (Sigma-Aldrich, C4790), glycyrrhetinic acid, ATP, Carbachol, thapsigargin (Invitrogen, T7459), native and n coelenterazines, BRD73954 (Tocris Bioscience, 18790528), BVT2733 (MedChemExpress, HY-18054), hydrocortisone cypionate (Fisher Scientific, 16402358), ^10^Panx1, GAP19, thapsigargin (Invitrogen, T7459), 2,5-di-tert-butylhydroquinone (BHQ), lanthanum (III) chloride heptahydrate (Sigma-Aldrich, 262072), probenecid (Invitrogen, P36400), digitonin (Sigma-Aldrich), Fura-2 AM.

### Synthesis of CBX derivatives

Materials, physical measurements and synthesis of CBX compounds and their chemical characterization and nuclear magnetic resonance and high-resolution mass spectra are described in Supplement Methods.

### Wound healing

Experiments were performed using HUVECs at passages 3-5. To perform a wound-healing assay, 100,000 cells/mL of HUVECs were plated on fibronectin-coated (1 µg/mL) FluoroDishes with glass cover bottoms (35 mm diameter) and incubated in full media for 2 days at 37°C and 5% CO_2_. When cells reached ∼90% confluency, nuclei were stained with Hoechst 33342 trihydrochloride, trihydrate (80 nM, #H1399) for 20 min at 37°C and 5% CO_2_.

A sterile 1000 µL micropipette tip was used to scratch the cell monolayer and create a wound. After the scratch, the cell monolayer was gently washed with full media to remove cell debris and any remaining floating cells. Then, the cells were treated before live-cell imaging, with 1 µM of thapsigargin (Invitrogen, T7459), 10 µM or 50 µM Carbenoxolone disodium salt (Sigma-Aldrich, C4790), 100 μM lanthanum (III) chloride heptahydrate (Sigma-Aldrich, 262072), or 500 μM probenecid (Invitrogen, P36400). The FluoroDishes were imaged in phase contrast and DAPI under culture conditions at 37°C and 5% CO_2_ using a Leica Dmi8 inverted microscope, 10x objective. The wound-healing assay was captured by taking pictures every 5 min.

Data Analysis: Before tracking cell nuclei during wound healing, the image positions covering the wound field of view were merged using the “Stitching” plugin (Preibisch et al., 2009). The stitched images from different conditions were then compared, and the condition in which wound closure occurred first was selected as the reference for the analysis duration. A region of interest (ROI) was defined on the frame where the two wound edges first came into contact, and this ROI was subsequently applied to all other conditions.

Cell migration was analysed by detecting the nuclei with TrackMate (estimated object diameter: 15–20 µm), followed by a Nearest-neighbour tracker (maxi. linking distance: 15–20 µm). Quality filters were implemented to remove non-cellular particles, and the values of speed were obtained directly from exporting the data from the software. The Mean square displacement (MSD) was calculated using MotilityLab (Wortel et al., 2021).

Collective cell migration experiments were also performed with erGAP3-ARPE-19 cells. In addition, 2D free single cell migration was performed by seeding 10,000 cells/mL in fibronectin coated FluoroDishes. Images were acquired every 40 s using the same microscope as above. The tracks were further analyzed using MotilityLab to calculate the mean square displacement (Wortel et al., 2021).

### Immunofluorescence and staining

HUVECs were grown on 12 mm coverslips (Th. Geier, 6080181) coated with fibronectin and to 70% confluency and treated with 50 µM fluorescent carbenoloxone (CBX-A) for 20 minutes at 37°C and 5% CO_2_. Then, cells were washed once with 1X DPBS, before fixation with 4% paraformaldehyde (Electron Microscopy Science, 15710) in 1X Dulbecco’s phosphate-buffered saline (DPBS) for 15 minutes, washed three times with 1X DPBS for 5 minutes each and stored in 1X DPBS at 4°C.

Antibody staining was performed on parafilm in a humid chamber. Blocking was done for 15 min at room temperature in blocking solution (1X DPBS, 2% bovine serum albumin [BSA, Sigma-Aldrich, A2153], 0.3 M glycine). Primary antibody solution was prepared in 1X saponin solution (10X saponin solution: 10X DPBS, 2% BSA, 0.5% saponin) (Saponin, CAS 8047-15-2, Merck Millipore, 558255) and staining was performed overnight at 4 °C. Next, the primary antibody staining was removed, and cells were washed once with 1X saponin solution, once with 1X DPBS, and a third time with 1X saponin solution before the secondary antibody staining. Secondary antibody staining was prepared in 1X saponin solution with the below-mentioned secondary antibodies as well as phalloidin and staining was done for 60 minutes at room temperature in the dark. Next, cells were washed three times with 1X DPBS and coverslips were mounted on microscope slides (Th. Geyer, 6052041) using Fluoromount-GTM (InvitrogenTM, ThermoFisher, 00-4958-02) and stored at 4 °C.

Rabbit polyclonal anti-NOGO-A antibody for ER (1:250, Novus Biologicals, NB100-56681SS), chicken anti-Pannexin1 antibody (1:200, Diatheva, ANT0027) were used as primary antibodies for immunostaining. Secondary antibodies used were goat anti-rabbit IgG (H+L) Alexa Fluor 647 (1:250, Thermofisher, A-21245) and CyTM3 AffiniPure F(ab’)2 Fragment donkey anti-chicken IgY (IgG) (H+L) (1:100, Jackson Immunoresearch, 703-166-155). F-actin was stained using Alexa FluorTM Plus 405 Phalloidin (1:100, Thermofisher, A30104).

Confocal images shown in Fig. 4A and B were acquired on a Zeiss Axio Examiner.Z1 with LSM 980 and Airyscan 2 (DFG code: INST 152/876-1 FUGG), equipped with a 63X, 1.40 NA Plan-APOCHROMAT Oil DIC objective (working distance 0.19 mm), and 405-nm, 488 nm, 561 nm and 639 nm continuous wave laser lines. Z-stack images were captured with a step-size of 0.25 µm and images were processed using the Zeiss Zen blue edition software. Images shown in Fig. 4D were acquired on a SP5 Leica confocal microscope equipped with a white laser and a 63X, 1.40 NA Plan-APOCHROMAT Oil DIC objective. Images were analyzed with ImageJ software using the JACoP plugin (Bolte & Cordelières, 2006).

### Single cell Ca^2+^ fluorescence imaging

ER Ca^2+^ was measured using erGAP3. For Ca^2+^ imaging experiments, cells expressing erGAP3 were seeded onto poly-L-lysine-coated 12 mm diameter glass coverslips at a density of ∼4×10^4^ cells. Imaging of erGAP3 was performed 1 day later using an upright fluorescence microscope (Axioplan, Zeiss) equipped with a 20X water-immersion objective (W-Achroplan, Zeiss; NA=0.5) and an AxioCam MRm camera (12 bits; Zeiss). The system was coupled to a xenon lamp with an excitation filter wheel containing 405/12DF and 470/25DF filters. GAP was excited at 470 and 405 nm sequentially at each excitation wavelength every 10 s and acquired at >515 nm emission (plus a 535DF35 filter). The system was run using Zeiss Axiovision software.

The dual measurements of ER and cytosolic Ca^2+^ were performed using erGAP3- and Fura-2, respectively. Cells were incubated with Fura-2 AM (4 μM) for 1h at 22°C in extracellular-like medium (EM) that contained 145 mM NaCl, 5 mM KCl, 1 mM MgCl_2_, 10 mM glucose, 10 mM Na-HEPES (pH 7.4) and 1 mM CaCl_2_. Fluorescence was recorded in a Nikon Diaphot equipped with an 20X objective (PlanApoUV; Olympus N.A. 0.7) and a filter wheel with the 403/12 DF y 470/25 DF filters and the DM500 dicroic mirror. The light above 520 nm (LP520) was collected with a Hamamatsu C4742-98 camera and the PCI 6.6 Simple Hamamatsu software.

Intact cells were recorded under continuous perfusion at 5–6 ml/min with EM. CBX or the mixture of ATP and carbachol were dissolved in EM and perfused during the time indicated. Each run ended with the perfusion of an ER Ca^2+^-depletion (Fmin) cocktail containing 10 μM SERCA inhibitor 2,5-ditert-butylbenzohydroquinone (BHQ), 0.5 mM EGTA and 100 μM ATP in EM medium without added CaCl_2_. All experiments were recorded at 22°C. Background was subtracted from the output images and pixel-to-pixel ratios were calculated using ImageJ software. The ratio R (F470/F405) was expressed as R/R_0_ or R/R_min_. R_0_ was computed as the mean of the ratios obtained during the first 5–10 frames of each experiment.

For high resolution experiments, erGAP3-ARPE-19 cells were grown on FluoroDishes (WPI (Freitext); FD35-100) until they reached 20% confluency and were then incubated with 1.5 µM Calbryte^TM^ 590 AM (Biomol GmbH, ABD-20700) for 30 min at 37°C, 5% CO_2_. Next, cells were washed once with 1X DPBS and DMEM F:12 full media was added. Cells were imaged immediately after staining in a Nikon Ax R confocal laser scanning microscope equipped with a Plan Apo λD 60x oil objective lens (NA 1.42). Images were acquired using bidirectional resonant scanning with a line averaging of 8, a resolution of 1024×1024 pixels and a resulting calibration of 0.09 µm/px. Acquired images were postprocessed using Denoise.ai of NIS elements 6.10. The time resolution of the recordings was about 1.2 seconds. A 60s baseline period was imaged prior to any treatment. Subsequently, carbenoxolone disodium salt (50 µM, Sigma-Aldrich, C4790) or thapsigargin (1 µM, Invitrogen, T7459) were added, and imaging was continued for an additional 90 s.

### ER Ca^2+^ measurements by luminescence

Cells seeded in 4-well plates were incubated for 10 min at 22 °C in a standard incubation medium with the following composition: 145 mM NaCl; 5 mM KCl; 1 mM MgCl_2_; 10 mM glucose; 10 mM sodium-HEPES, pH 7.4; 0.5 mM EGTA and 10 μM BHQ. Apo-aequorin was then reconstituted by incubation with 1 µM coelenterazine *n* added in the same medium during 1-2 hours prior to measurements. The cells were placed in a chamber of a built-in-luminometer (Cairn Research, UK) and perfused with the same standard medium with 1 mM CaCl_2_ at 5 ml/min.

### ATPase activity

For experiments in permeabilized cells, EM containing 1 mM CaCl_2_ was perfused for 2 min. Then, cell plasma membrane was permeabilized by perfusing an intracellular-like medium (IM) containing no Ca^2+^ (140 mM KCl, 1 mM KH_2_PO_4_/K_2_HPO_4_, 1 mM MgCl_2_, 0.2 mM EGTA, 1 mM Na-pyruvate, 2 mM Na succinate, 1 mM ATP/MgCl_2_, 20 mM K-HEPES pH 7.2) and 20 µM digitonin for 1 min. Cells were then perfused with IM containing 200 nM free Ca^2+^ (buffered according to MAXCHELATOR (Bers et al., 2010). Unless indicated otherwise, all the reagents added were dissolved in IM. Recordings were collected every second. At the end of each experiment, cells were lysed with 100 μM digitonin in a 10 mM CaCl_2_ solution to release all the aequorin luminescence. The calibration of luminescence signal into [Ca^2+^] requires calculating the L/L_t_ ratio, where L is the luminescence emission (in cps) at a given time, and L_t_ is the sum of all the counts remaining at that time (Rodriguez-Prados et al., 2025).

For the experiments exploring the effects of inhibitors acting against alternative targets of CBX, 100 µL stably-expressing IgGAP1-Hela cells (10^4^ cells) were seeded in 96-well microplates. Next, cells were reconstituted by incubation with 1 µM native coelenterazine for 1h prior to measurements in a TECAN plate reader. Cells were permeabilized in IM without ATP containing 200 nM Ca^2+^. ATPase activity was triggered by injection of 1 mM ATP. Acquisition was performed every 1s. At the end of each run, a calibration cocktail was injected.

### Molecular docking

The structure of the SERCA1a protein (PDB-code: 3N5K) was co-crystallised with the inhibitor thapsigargin. This binding site was prepared for a docking experiment, i.e. all aminoacids were checked for completeness, hydrogens added, protonation states assigned. This served as input for the docking algorithm SurflexDock v.2.51 from BioPharmics IT (Sonoma County, CA, USA) as implemented in the software package SYBYL-X1.3 (Tripos, St. Louis, MO, USA) to place the ligands.

### Statistical analysis

Results are shown as mean ± S.E.M., as indicated. Raw data were initially processed using Origin 2024 software (Origin Lab™), and all statistical analyses were conducted with GraphPad Prism 10 (San Diego, CA, USA). After confirming the normal distribution of data by the Shapiro-Wilk test, statistical significance between two groups was determined by the Student’s *t*-test for parametric data, while comparisons involving more than two groups were performed using one-way ANOVA followed by Tukey’s post-hoc test. Statistical significance was determined for probability (*p*) values below 0.05 (*), 0.01 (**), 0.001 (***) or 0.0001(****). Sample sizes (n) correspond to independent cell wells from at least three separate experimental days. Representative experiments are shown as individual traces.

## ACKNOWLEDGEMENTS

The authors thank Jesús Fernández, Carla Rodríguez, Iris Fernández and Johan M. Kux for expert technical help. We also thank Dr. A. Guerrero (CINVESTAV, Mexico) for his suggestions and Dra. A. M. Mata (U. Extremadura, Spain).

## AUTHOR CONTRIBUTIONS

All authors participated in analysis, discussion, and interpretation of data, revised the article, and gave final approval.

Conceptualization: P.J.S. and M.T.A. Methodology: P.J.S. and M.T.A. Software: S.M.P. and P.J.S. Formal analysis: C.S.-R., B.C., S.M.P., M.R.A., J.R.-R. Investigation: C.S.-R., B.C., S.M.P. M.R.A. C.C., J.R.-R. Resources: J.L.C-M, A.J.-S. Data curation: C.S.-R., B.C., S.M.P. M.R.A., U.U. Writing—original draft: P.J.S. and M.T.A. Writing—review and editing: C.S.-R., B.C., S.M.P. M.R.A., J.L.C-M, A.J.-S., M.L.G.-G., U.U., T.S., J.R.-R. Visualization: C.S.-R., B.C., S.M.P., M.R.A., U.U., P.J.S., and M.T.A. Supervision: A.J.-S., P.J.S. and M.T.A. Project administration: A.J.-S., P.J.S. and M.T.A. Funding acquisition: A.J.-S., P.J.S. and M.T.A.

## DATA AND MATERIALS AVAILABILITY

The data underlying this article will be shared on reasonable request to the corresponding author.

## Supplement Figure Legends

**Supplement Figure 1.**
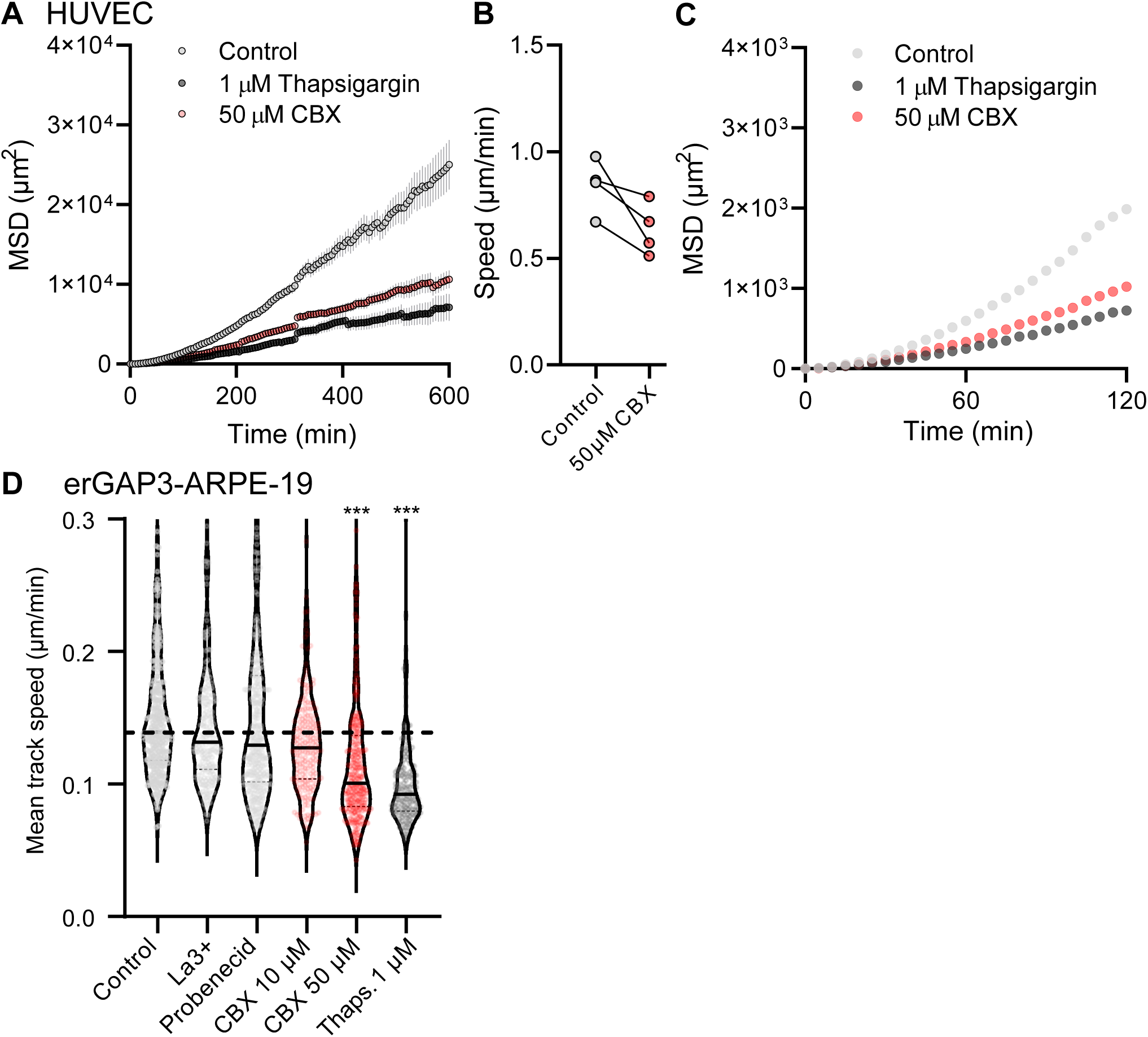
Collective cell migration depends on ER Ca^2+^. **(A)** Graph showing the mean square displacement (MSD) in HUVECs undergoing collective migration. **(B)** Pooled data shows the median instantaneous speed in four independent experiments, showing the effect of CBX. **(C)** Similar graph to the one shown in A but highlighting the first 2 h of collective migration. **(D)** Graph showing the speed of migration in erGAP3-ARPE-19 cells in one of three representative experiments (n=3; n>200 cells per condition, per experiment). The line denotes the median. ***p<0.01 Kruskal-Wallis test compared to Control condition.

**Supplement Figure 2.**
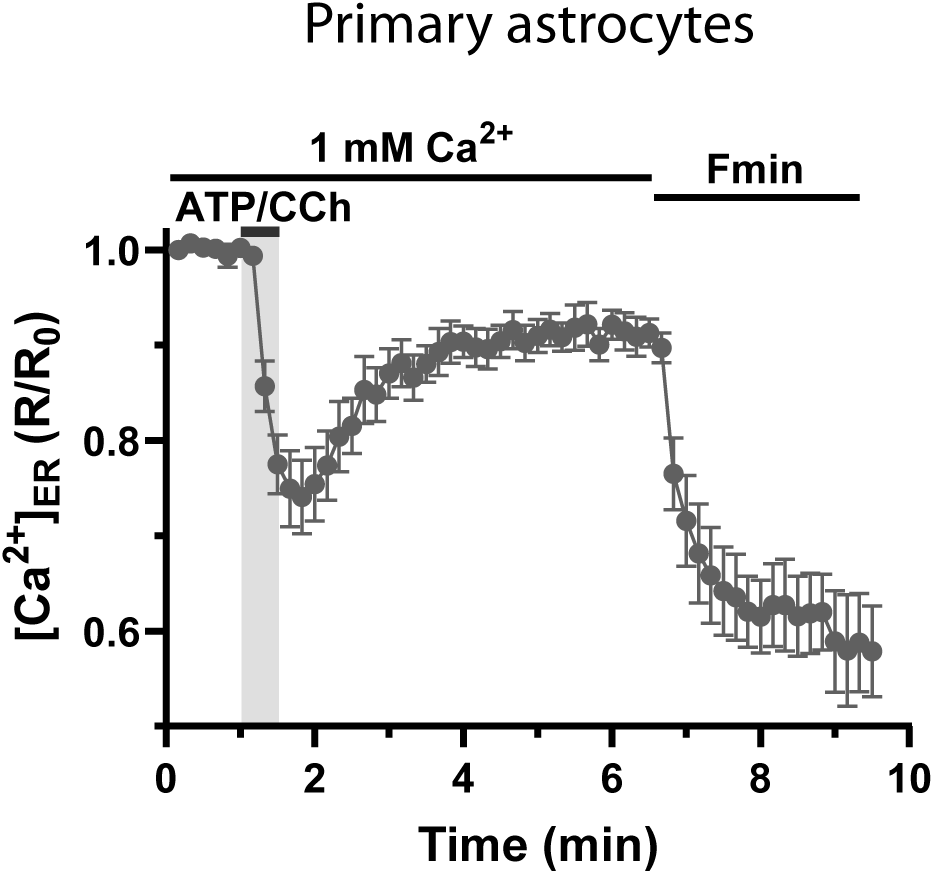
CBX empties the ER of Ca^2+^ with kinetics different to that by activation of IP_3_Rs. Ca^2+^ release from the ER of mouse cortical astrocytes isolated from erGAP3 transgenic mice stimulated with ATP+Carbachol (ATP/CCh; 100 µM each), two IP_3_-generating agonists. [Ca^2+^]_ER_ is represented as F470/F405 ratio (R) normalized to R_0_ (R/R_0_). The trace is the mean ± SEM of 6 cells.

**Supplement Figure 3.**
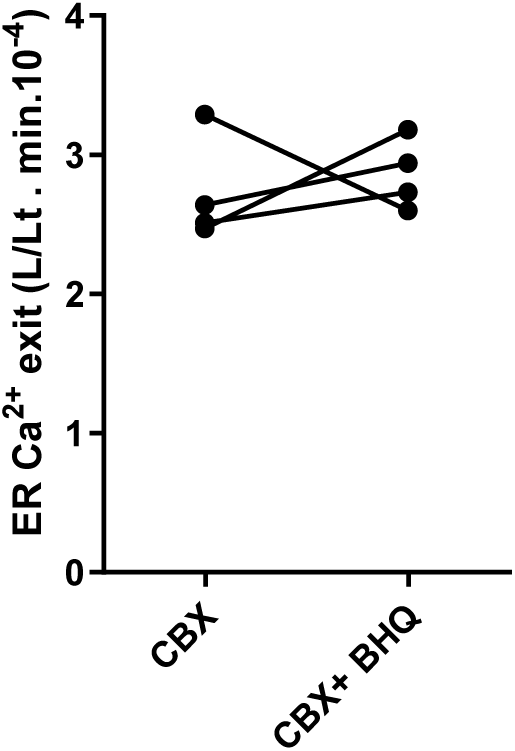
Summary data (mean ± SEM; n=4) from the traces shown in Fig. 3H. Student′s *t* test was applied; no statistically significant differences were found.

**Supplement Figure 4.**
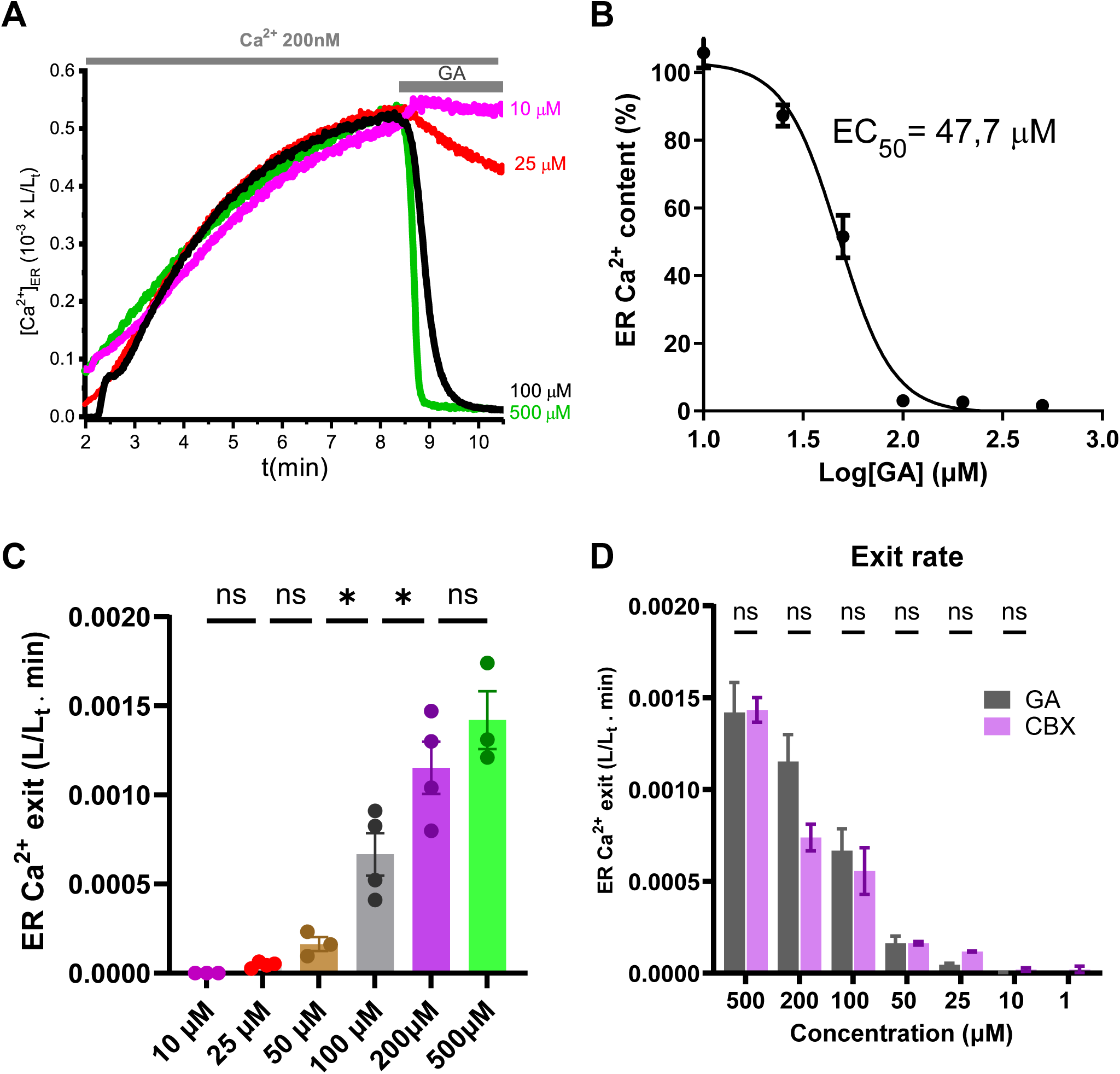
The gap-junction inhibitor glycyrrhetinic acid passively depletes Ca^2+^ from the ER. **(A)** Effect of various concentrations of glycyrrhetinic acid (GA) on the ER Ca^2+^ in digitonin-permeabilized er(Ig)GAP1-HeLa cells. Other experimental details as in Fig. 3. **(B)** Dose-response of glycyrrhetinic acid on the ER Ca^2+^ content (represented as percentage), measured at min 3 upon GA addition. Data are represented as mean ± SEM (n = 3-4). **(C)** Concentration-dependent response of GA on the exit of Ca^2+^ from the ER represented as initial rate (L/Lt·min). Data are represented as mean ± SEM (n = 3–4). **(D)** Comparison of the effects of CBX and GA on the exit of Ca^2+^ from the ER represented as initial rate (L/Lt·min). Summary quantification (mean±SEM) of 3-4 coverslips (from two independent days). Student′s *t* test was applied, and no statistically significant differences were found.

**Supplement Figure 5.**
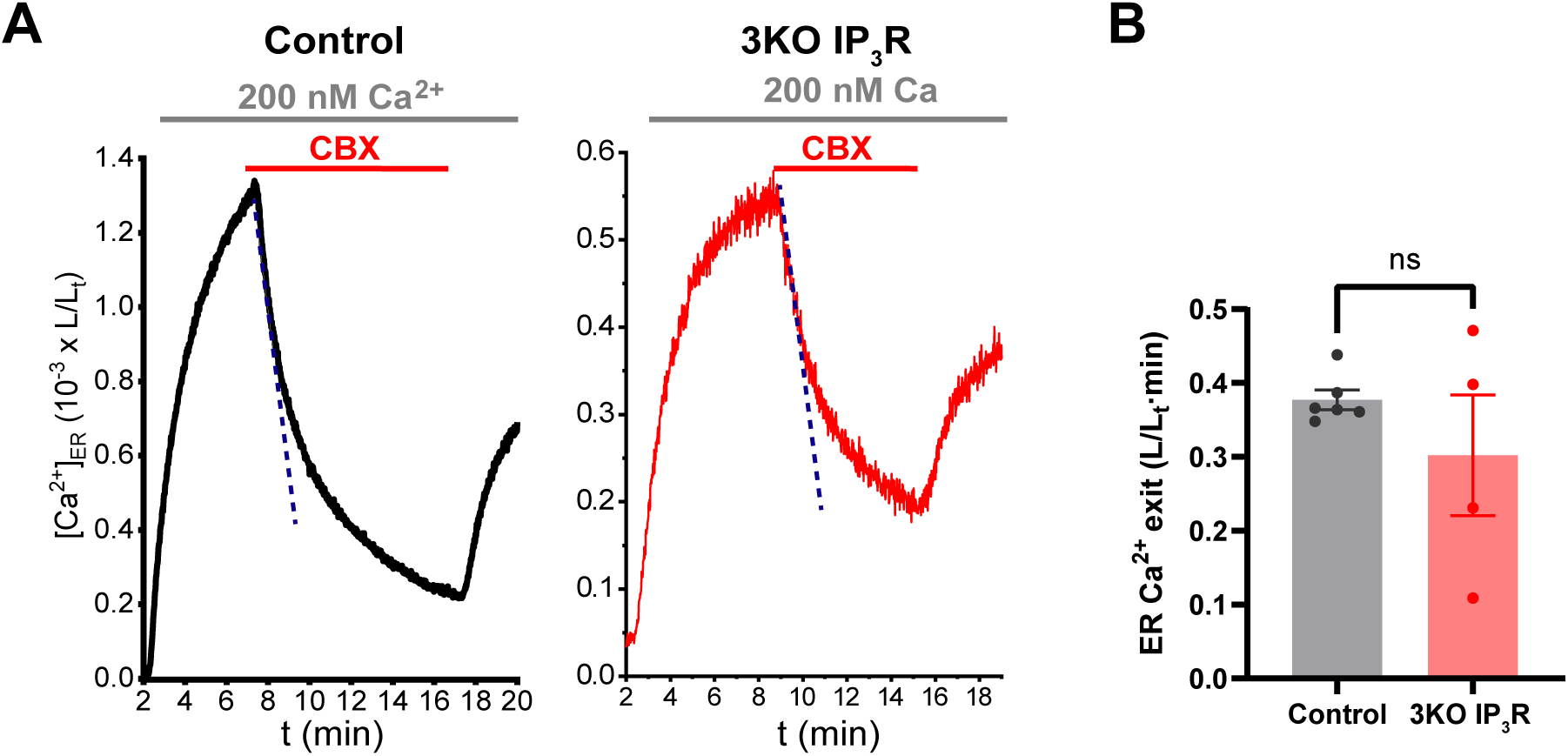
IP_3_R is not required for the effect of CBX. (**A)** Response to CBX (100 µM) in wt (Control) and IP_3_R-null (3KO IP_3_R) HEK293 cells. (**B)** Pooled data (n=4-6) of similar experiments to **A**. Each point represents ER Ca^2+^ exit (L/Lt·min) normalized for the ER Ca^2+^ at min 8. Data are represented as mean ± SEM. Student′s *t* test was applied and no statistically significant differences were found.

**Supplement Figure 6.**
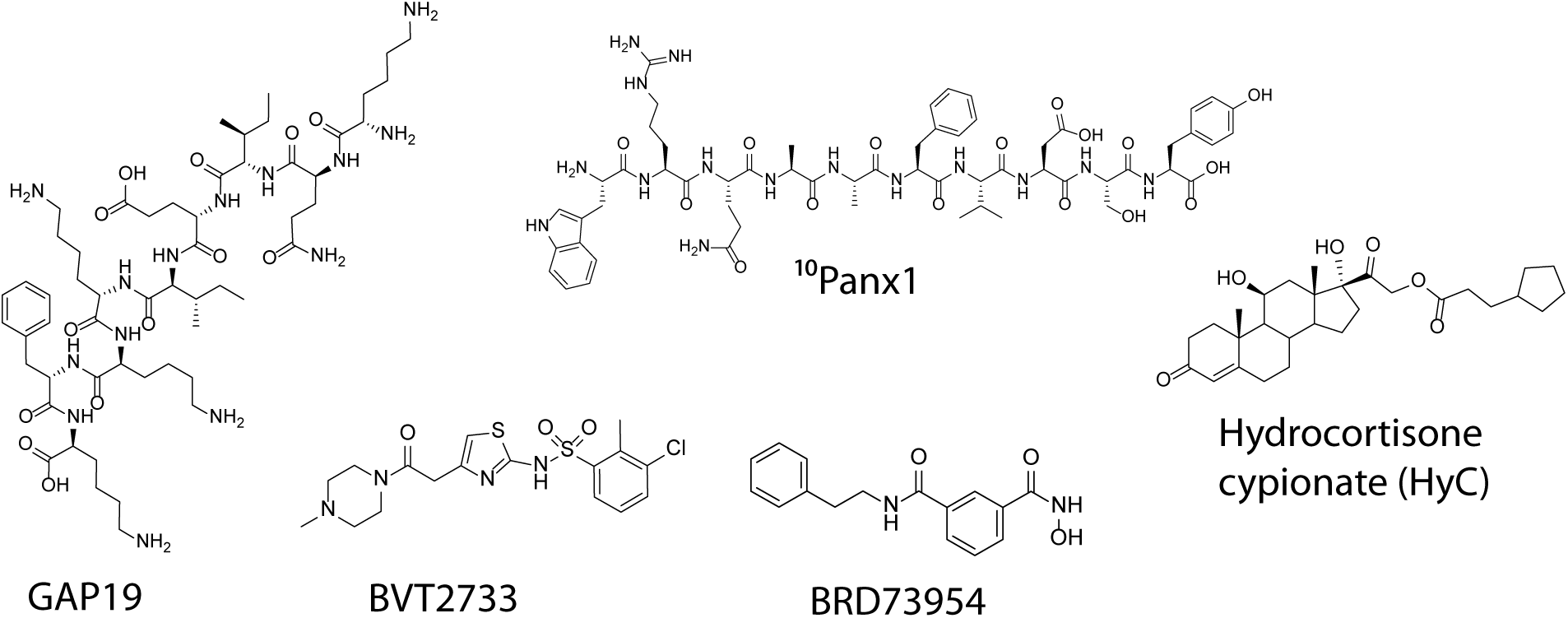
Chemical structure of inhibitors that act on alternative targets of CBX.

**Supplement Figure 7.**
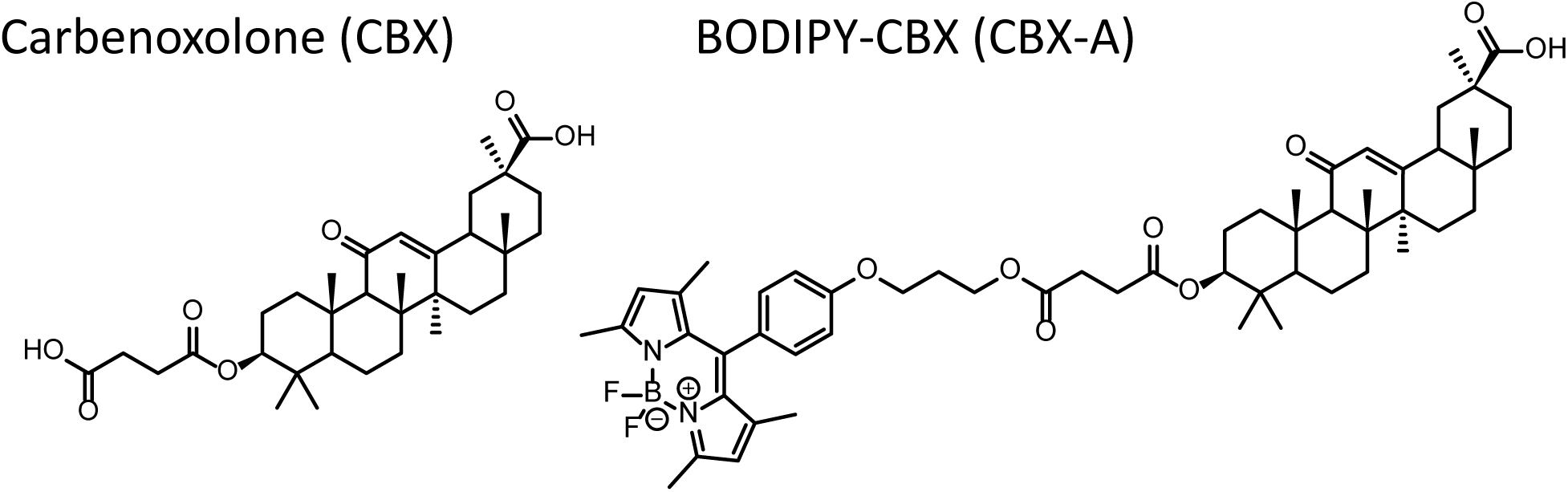
Chemical structure of carbenoxolone (CBX) and its fluorescent derivative CBX-A.

## ABBREVIATIONS

GFP: green fluorescent protein
GAP: GFP-Aequorin Protein
ER: endoplasmic reticulum
SERCA: sarco/endoplasmic reticulum Ca^2+^ ATPase
[Ca^2+^]_C_: cytosolic free Ca^2+^ concentration
[Ca^2+^]_ER_: Ca^2+^ concentration inside ER. TBH, 2,5-di-tert-butylhydroquinone
IP_3_: inositol 1,4,5, trisphosphate
CCh: carbachol.

