## Supplement Methods for "Carbenoxolone disrupts cell migration by inhibiting the SERCA pump"

1. Experimental procedures S2
1.1. Materials and physical measurements S2
1.2. Compounds synthesis and chemical characterization S2
1.3. Nuclear Magnetic Resonance Spectra S4
1.4. High-resolution mass spectra S8
1.5. References S8

### 1. Experimental Procedures

#### 1.1. Materials and physical measurements.

All starting materials were purchased from Aldrich and used without further purification. Solvents were dried by standard methods or distilled prior to use. Reactions were monitored by TLC on pre-coated silica gel plates (ALUGRAM SIL G/UV254) and revealed by exposure to a UV254 lamp. Commercially available starting materials, and solvents were used as supplied. $^{1}H$ and $^{13}C$ NMR spectra were recorded at room temperature on a 400 MHz Bruker and 300 MHz Jeol unity spectrometer. Chemical shifts (ppm) are relative to $(CH_3)_4Si$. High resolution mass spectrometry (ESI-TOF) was obtained by using an Agilent Technologies 6530 Accurate-Mass Q-TOF LC/MS equipment.

#### 1.2. Compounds synthesis and chemical characterization

Scheme S1 describes the synthetic methodology followed to prepare the **Carbenoxolone derivative**.

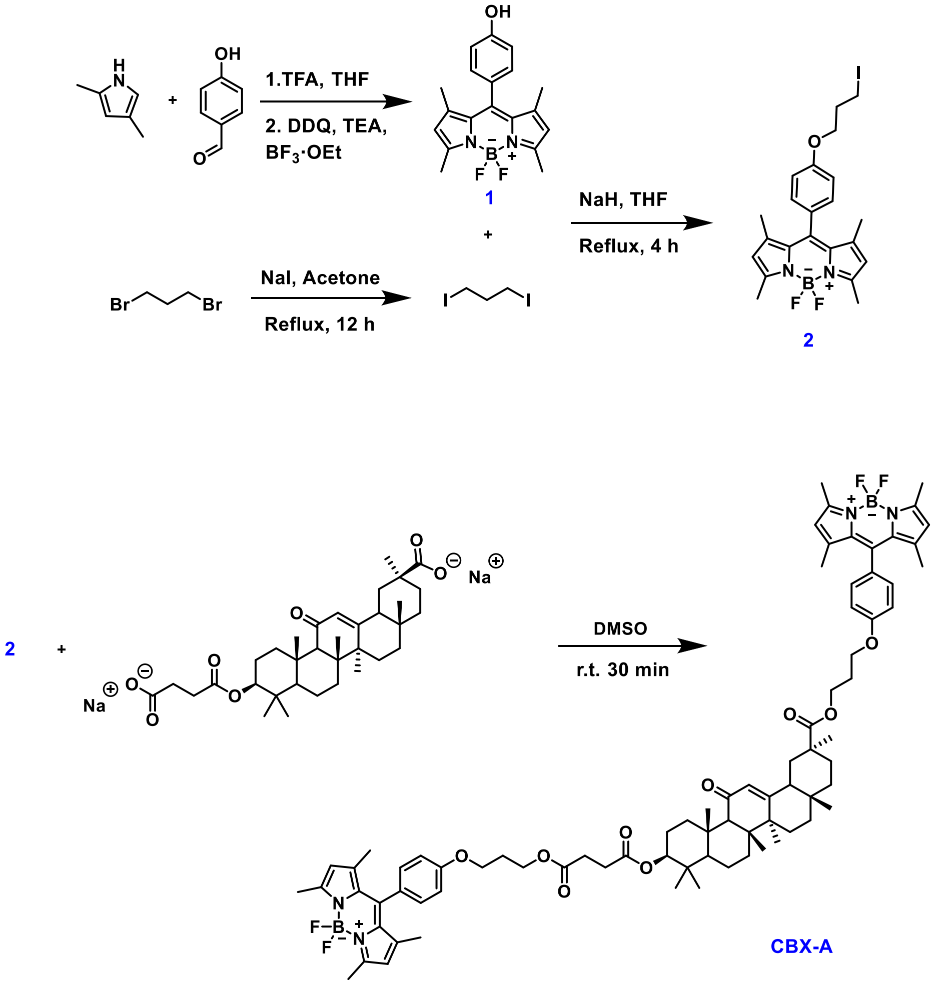

**Scheme S1**. General synthetic methodology to obtain **CBX** derivatives. See the text for synthetic details and chemical characterization.

#### Synthesis of BODIPY derivative

Synthesis of 4-(5,5-difluoro-1,3,7,9-tetramethyl-5*H*-4λ$^4$,5λ$^4$-dipyrrolo[1,2-*c*:2',1'-*f*][1,3,2]diazaborinin-10-yl)phenol **(1)** [1]2,4-Dimethylpyrrole (0.5 g, 5.2554 mmol) and 4-Hydroxybenzaldehyde (0.320 g, 2.627 mmol) were dissolved in anhydrous etrahydrofurane(THF) (20 mL). Then several drops of trifluoroacetic acid were added and the mixture was stirred at room temperature (25ºC) for 12 h. Later, tetrachloro-1,4-benzoquinone (0.625 g, 2.5418 mmol) dissolved in 20 mL of THF was added dropwise. The mixture was stirred at 25ºC for 4 h. After the addition of (diisopropylethylamine (DIPEA) (5.56 mL, 39.4 mmol), the mixture was cooled in ice-water bath and $BF_3{\cdot}Et_2O$ (5 mL, 46.5 mmol) was added dropwise. The mixture was stirred for 2 h. The solvent was removed under reduced pressure and the residue was dissolved with $CH_2Cl_2$ (50 mL). This organic phase was washed with a saturated aqueous NaCl solution (2 × 100 mL). The organic portion was collected and dried over anhydrous $Na_2SO_4$. Removal of the sodium sulfate by filtration and evaporation of the solvent under reduced pressure led to a crude product that was purified with column chromatography (silica gel, ethyl acetate/hexane, 5:95 v/v). Deep red solid powder was obtained (0.232 g; Yield:26%). $^1$H NMR (300 MHz, $CDCl_3$) δ (ppm): 7.11 (d, *J* = 8.5 Hz, 2H), 6.94 (d, *J* = 8.5 Hz, 2H), 5.98 (s, 2H), 2.55 (s, 6H), 1.44 (s, 6H).

Synthesis of 5,5-difluoro-10-(4-(3-iodopropoxy)phenyl)-1,3,7,9-tetramethyl-5*H*-4λ$^4$,5λ$^4$-dipyrrolo[1,2-*c*:2',1'-*f*][1,3,2]diazaborinine **(2)**

Compound **1** (100 mg, 0.2939 mmol) and sodium hydride (suspension 60% in paraffin oil) (53 mg, 1.3251 mmol) were dissolved in anhydrous THF and the mixture was stirred at 25ºC for 2h under inert argon atmosphere. Then, 1,3-diiodopropane was added to the solution and heated at reflux for 4h. Finished the reaction, solvent was removed under reduced pressure and the crude product was supported on celite and purified by column chromatography over silica gel with hexane/ethyl acetate as eluent in a 98:2 to 90:10 gradient to afford 60 mg of **2** (40% yield) as a red solid. $^1$H NMR (300 MHz, $CDCl_3$), δ (ppm): 7.17 (d, *J* = 8.6 Hz, 2H), 7.01 (d, *J* = 8.6 Hz, 2H), 5.98 (s, 2H), 4.09 (t, *J* = 5.9 Hz, 2H), 3.40 (t, *J* = 6.7 Hz, 2H), 2.55 (s, 6H), 2.38 – 2.27 (m, p, 2H), 1.43 (s, 6H).

#### Synthesis of CBX-A conjugate

Carbenoxolone disodium salt (72.58 mg, 0.1180 mmol) and compound **2** (20 mg, 0.03935 mmol) were dissolved in anhydrous dimethyl sulfoxide (DMSO) and the mixture was stirred under nitrogen atmosphere at 25ºC for 30 minutes. The product was extracted three times with a mixture DCM/$H_2O$ and the organic phase was collected and dried over anhydrous $Na_2SO_4$.. Removal of the sodium sulfate by filtration and evaporation of the solvent under reduced pressure led to a brownish colored oil. The crude product was purified by column chromatography (silica gel, Hexane:AcOEt 98:2 to 90:10 gradient). And a deep three orange solid powder was obtained (10.47 mg; Yield: 22%).

[1]H NMR (400 MHz, $CDCl_3$), δ (ppm): 7.20 – 7.12 (m, 4H), 7.02 – 6.97 (m, 4H), 5.97 (d, *J* = 1.7 Hz, 4H), 5.66 (s, 1H), 4.54 (dd, *J* = 11.7, 4.7 Hz, 1H), 4.33 (t, *J* = 6.4 Hz, 4H), 4.10 (td, *J* = 6.1, 2.7 Hz, 4H), 2.83 – 2.74 (m, 1H), 2.65 (s, 3H), 2.55 (s, 12H), 2.35 – 2.32 (m, 1H), 2.17 (q, *J* = 6.3 Hz, 6H ), 1.43 (s, 8H), 1.42 (s, 6H), 1.36 (s, 2H), 1.35 (s, 3H), 1.26 (s, 3H), 1.15 (s, 6H), 1.12 (s, 4H), 0.88 (s, 6H), 0.79 (s, 3H).

[13]C NMR (100 MHz, $CDCl_3$), δ (ppm): 205.5, 200.3, 200.1, 195.5, 192.2, 187.1, 176.5, 172.5, 172.1, 169.4, 169.3, 159.5, 159.5, 157.6, 155.4, 155.4, 143.3, 143.3, 142.0, 132.00, 129.4, 129.4, 128.7, 128.6, 127.4, 127.3, 121.3, 115.2, 81.2, 78.9, , 64.7, 64.7, 62.0, 61.9, 61.8, 61.5, 55.2, 55.1, 48.6, 45.6, 44.2, 44.2, 43.4, 43.3, 41.2, 41.2, 39.3, 39.3, 38.9, 38.3, 38.2, 37.9, 37.2, 37.1, 32.9, 32.8, 32.0, 32.0, 31.3, 29.8, 29.8, 29.7, 29.5, 29.4, 28.9, 28.8, 28.7, 28.5, 28.5, 28.2, 28.2, 26.6, 26.6, 26.5, 26.5, 23.7, 23.5, 23.4, 22.8, 22.8, 18.8, 17.5, 16.8, 16.5, 16.5, 14.8, 14.7.

ESI-MS: m/z calc. for $C_{78}H_{96}B_2F_4N_4O_9^+$: 1330.7299; found: 1348.7741 $[M+F]^+$.

#### 1.3. Nuclear Magnetic Resonance Spectra

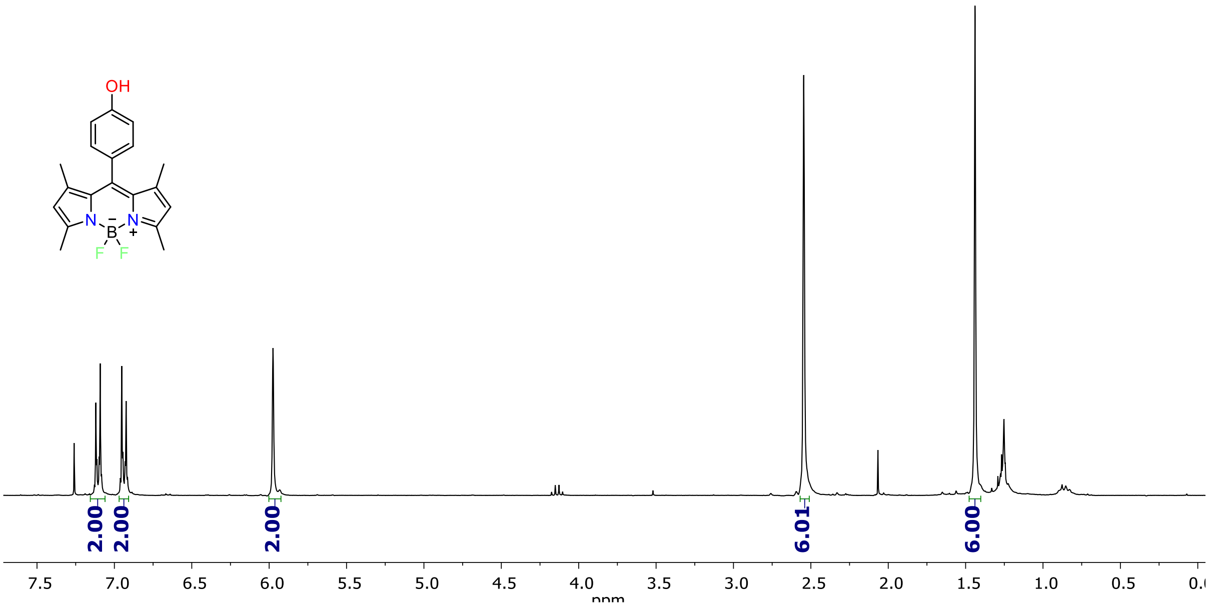


**Figure S1**. [1]H NMR spectrum of **1** in $CDCl_3$ (300 MHz).

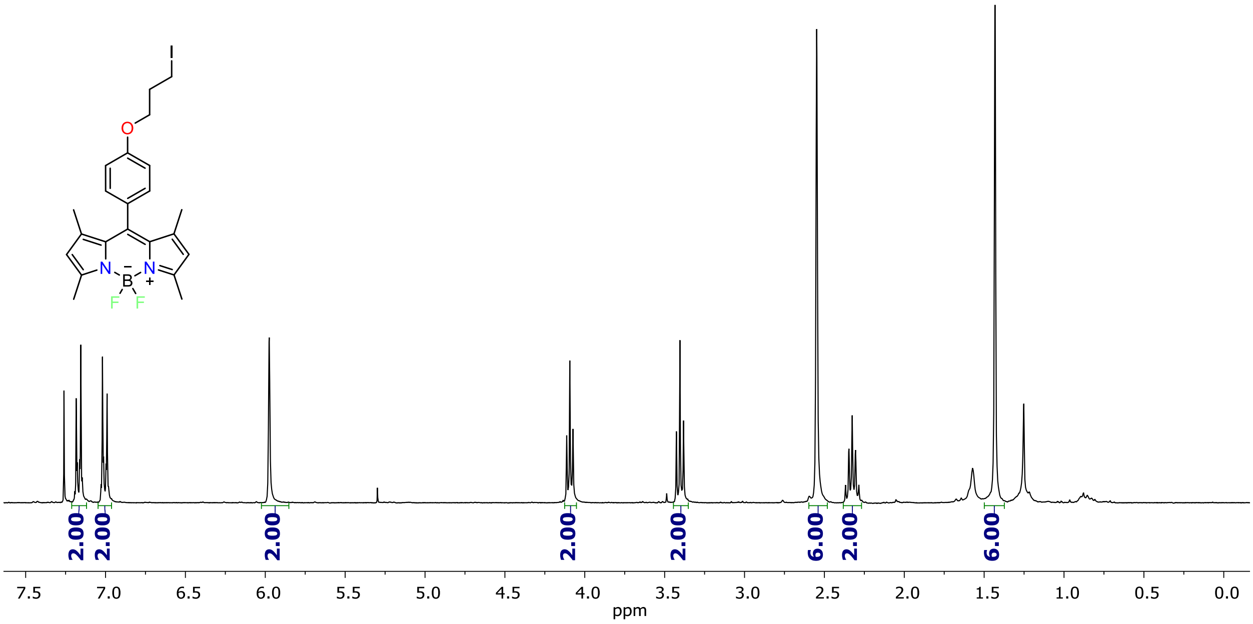


**Figure S2**. $^{1}H$ NMR spectrum of **2** in $CDCl_3$ (300 MHz).

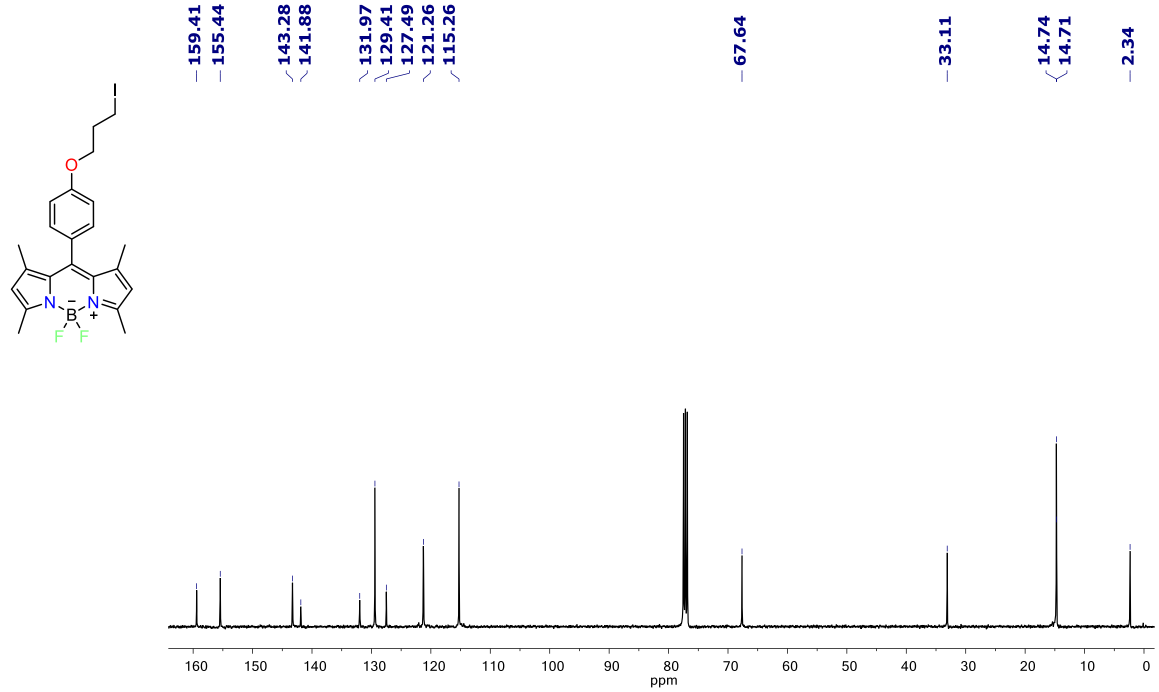


**Figure S3**. $^{13}C$ NMR spectrum of **2** in $CDCl_3$ (100 MHz).

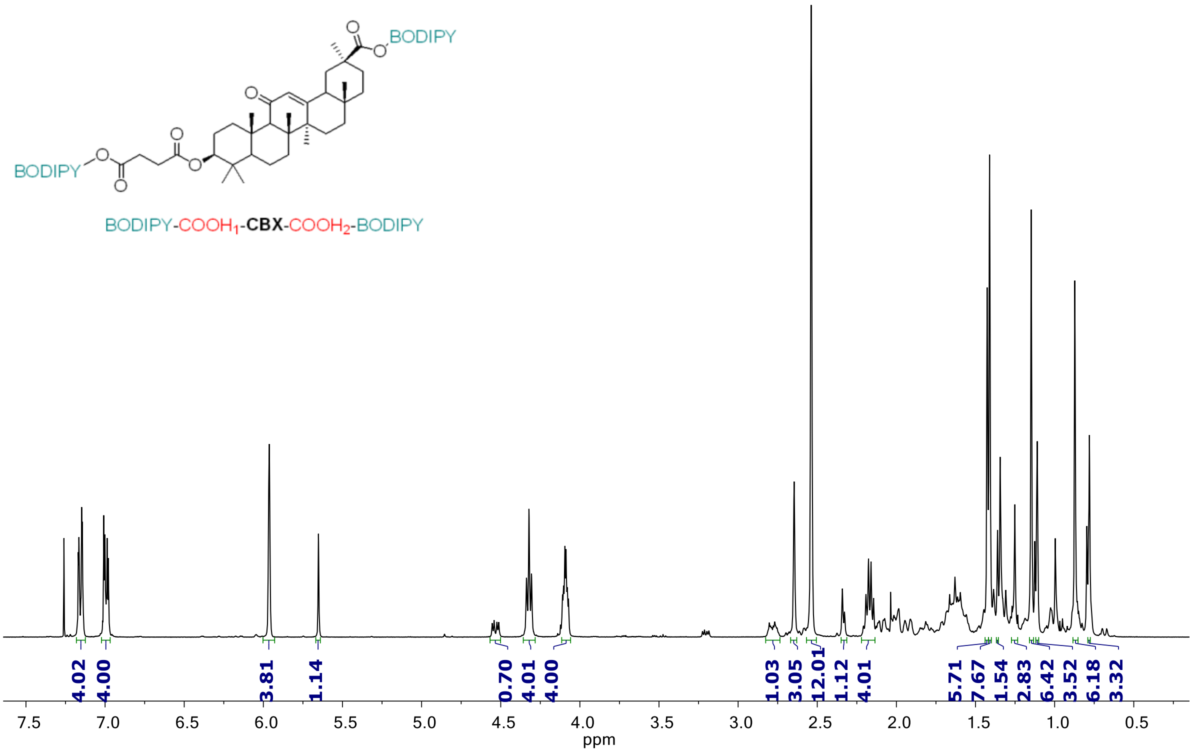


**Figure S4** . [1]H NMR spectrum of **CBX-A (BODIPY-COOH₁-CBX-COOH₂-BODIPY)** in $CDCl_3$ (400 MHz).

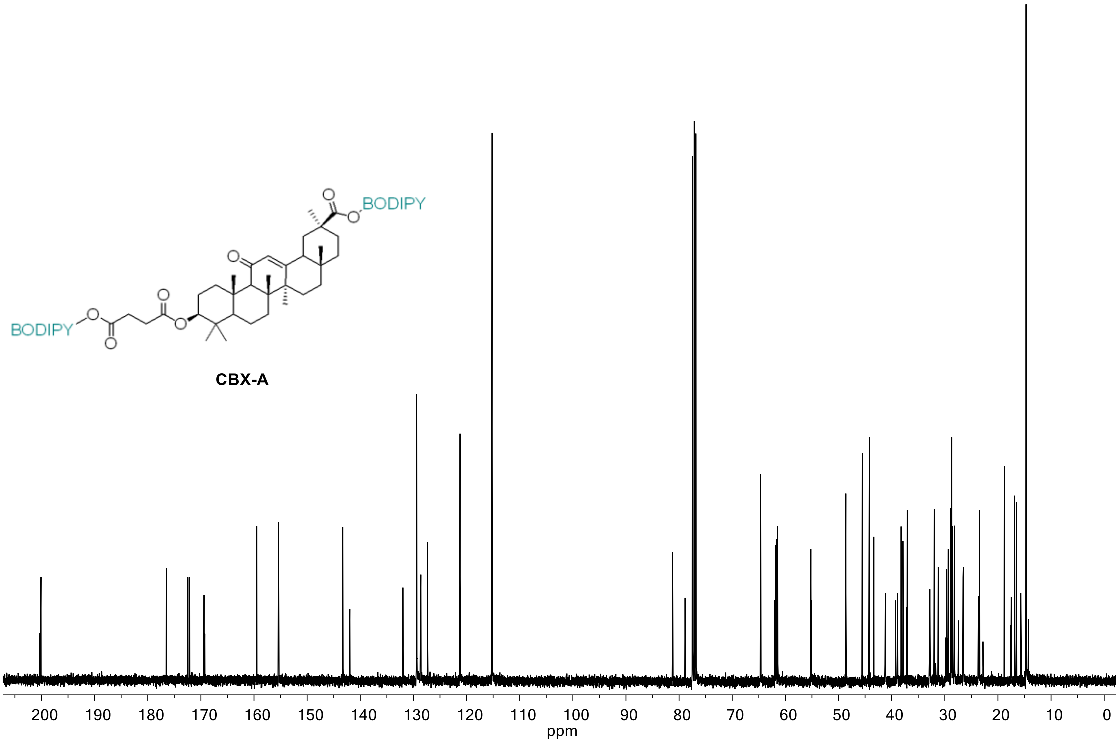


**Figure S5** . [13]C NMR spectrum of **CBX-A** in $CDCl_3$ (100 MHz).

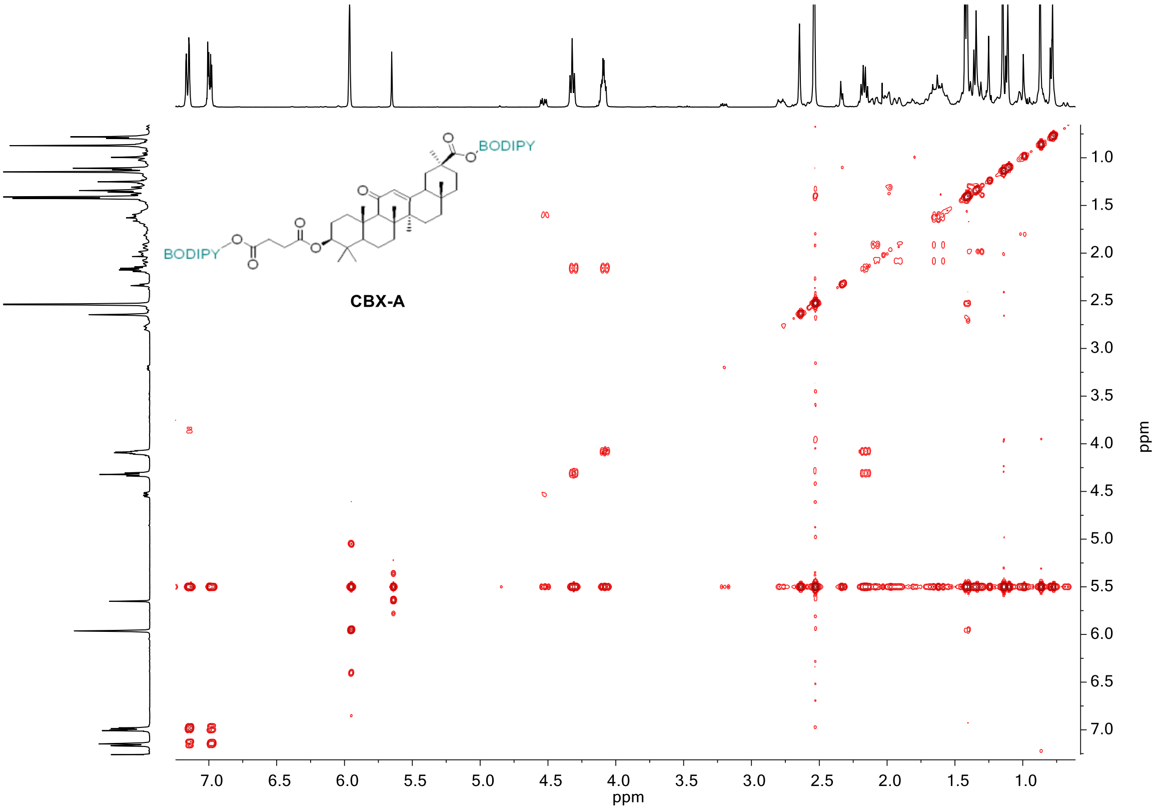


**Figure S6** . COSY experiment of **CBX-A** in $CDCl_3$.

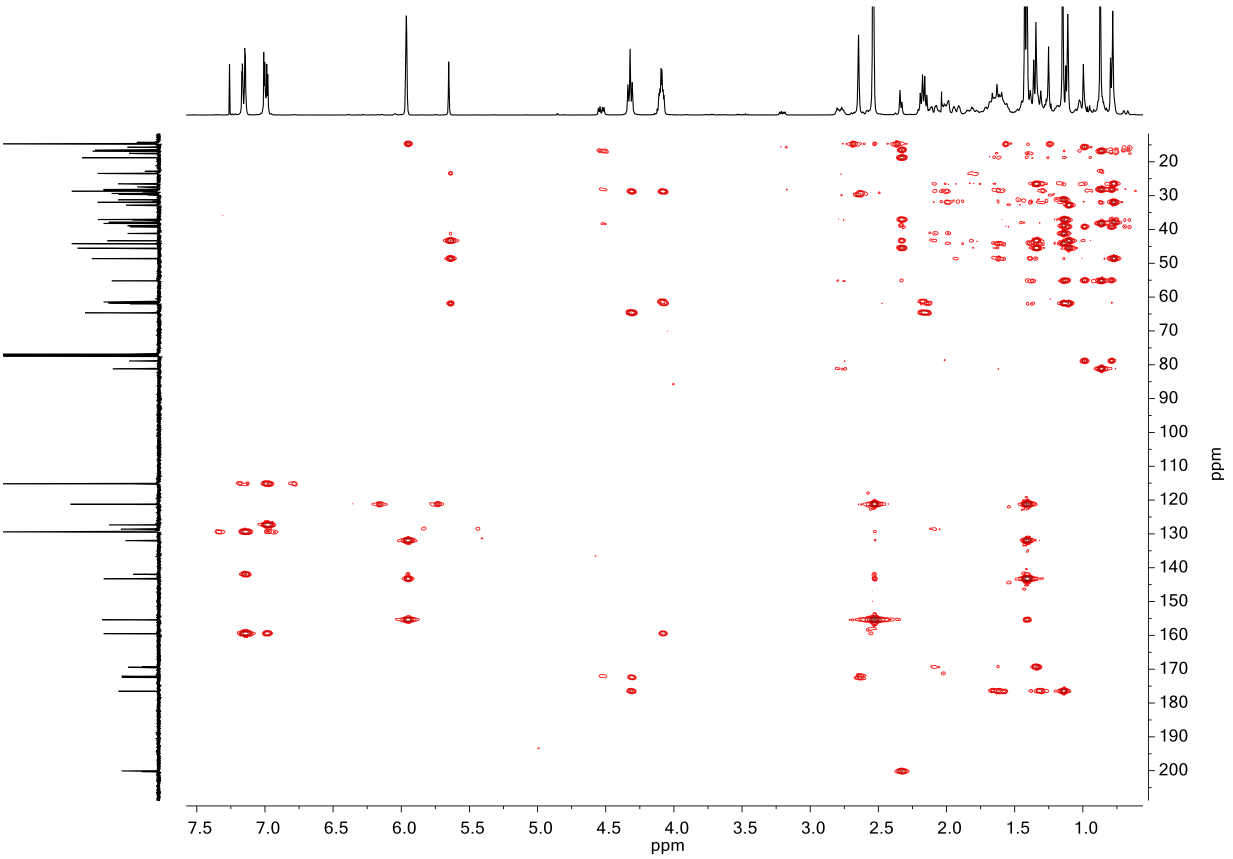


**Figure S7** . HMBC experiment of **CBX-A** in $CDCl_3$.

#### 1.4. High-resolution mass spectra

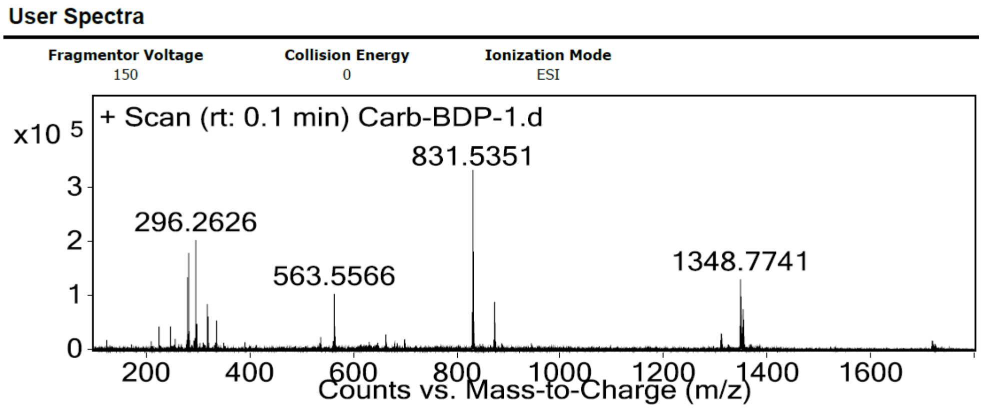


**Peak List**

| *m/z* | z | Abund |
|---|---|---|
| 280.2674 | 1 | 135846.18 |
| 282.2831 | 1 | 181872.68 |
| 296.2626 | 1 | 206513.36 |
| 318.2446 | 1 | 84650.82 |
| 563.5566 | 1 | 105069.64 |
| 831.5351 | 1 | 334327.42 |
| 832.5383 | 1 | 182322.68 |
| 873.5231 | 1 | 88555.09 |
| 1348.7741 | 1 | 130952.58 |
| 1349.7759 | 1 | 99231.85 |

**Figure S8 .** High-resolution mass spectrum (ESI-MS) of **CBX-A.**
